# Conformist social learning leads to self-organised prevention against adverse bias in risky decision making

**DOI:** 10.1101/2021.02.22.432286

**Authors:** Wataru Toyokawa, Wolfgang Gaissmaier

## Abstract

Given the ubiquity of potentially adverse behavioural bias owing to myopic trial-and-error learning, it seems paradoxical that improvements in decision-making performance through conformist social learning, a process widely considered to be bias amplification, still prevail in animal collective behaviour. Here we show, through model analyses and large-scale interactive behavioural experiments with 585 human subjects, that conformist influence can indeed promote favourable risk taking in repeated experience-based decision making, even though many individuals are systematically biased towards adverse risk aversion. Although strong positive feedback conferred by copying the majority’s behaviour could result in unfavourable informational cascades, our differential equation model of collective behavioural dynamics identified a key role for increasing exploration by negative feedback arising when a weak minority influence undermines the inherent behavioural bias. This ‘collective behavioural rescue’, emerging through coordination of positive and negative feedback, highlights a benefit of collective learning in a broader range of environmental conditions than previously assumed and resolves the ostensible paradox of adaptive collective behavioural flexibility under conformist influences.

## Introduction

Although the potential cost of strong conformist social influences in decision making has been well recognised (***Boyd and Richerson, 1985***; ***Kandler and Laland, 2013***; ***Kendal et al., 2005***; ***King and Cowlishaw, 2007***; ***Raafat et al., 2009***; ***Giraldeau et al., 2002***; ***Toyokawa et al., 2019***), collective learning through a tendency to engage in copying others’ behaviour has been considered widely adaptive in a range of environmental conditions (***Rendell et al., 2010***; ***Kameda and Nakanishi, 2002***; ***Toyokawa et al., 2014***, ***2019***; ***Miu et al., 2018***). One rationale behind this optimistic view might come from the assumption that individuals tend to prefer a behavioural option that provides larger net benefits in utility over those providing lower outcomes. Therefore, even though uncertainty makes individual decision making fallible, statistical filtering through informational pooling may be able to reduce uncertainty by cancelling out such noise. As a consequence, the group as a whole can boost its probability of choosing the benefit-maximising behaviour.

However, as numerous examples in behavioural science have documented, humans and many other non-human animals may not always choose or prefer a benefit-maximising behaviour over other behavioural alternatives. Instead, trial-and-error learning and adaptive exploration can constrain animals to be risk averse, which may prevent individuals from exploiting a high-risk high-return be-havioural option (***Hertwig and Erev, 2009***; ***Yechiam et al., 2006***; ***Weber, 2006***; ***March, 1996***). Although risk sensitivity can be adaptive in many ecological circumstances (***Real and Caraco, 1986***; ***McNamara and Houston, 1992***; ***Yoshimura and Clark, 1991***), such a risk-taking bias constrained by the fundamental nature of learning may function independently from the adaptive risk perception (***Frey et al., 2017***), potentially preventing adaptive risk taking. Therefore, the ostensible prerequisite of collective intelligence, that is, that individuals have a higher chance of exhibiting a more profitable behaviour than choosing a less profitable alternative, may not always hold in reality. Previous studies have suggested that individual decision making is biased further by social influences (***Chung et al., 2015***; ***Bault et al., 2011***; ***Suzuki et al., 2016***; ***Shupp and Williams, 2008***; ***Jouini et al., 2013***; ***Moussaïd et al., 2015***), which may nourish the spread of risk-related behavioural biases (***Dussutour et al., 2005***).

Given that behavioural biases are ubiquitous and learning animals rarely escape from them, it may seem that conformist social influences may often lead to suboptimal herding or collective illusion through recursive amplification of the majority influence (i.e., positive feedback; ***Denrell and Le Mens, 2007***, ***2016***). However, the collective improvement of decision accuracy has been widely observed in the real world in a wide range of taxa (***Sasaki and Biro, 2017***; ***Seeley et al., 1991***; ***Alem et al., 2016***; ***Sumpter, 2006***; ***Camazine et al., 2001***), including the very animals known to exhibit the learnt risk-taking biases that may incur large opportunity costs in particular circumstances, such as bumblebees (***Real, 1981***; ***Real et al., 1982***), honeybees (***Drezner-Levy and Shafir, 2007***), and pigeons (***Ludvig et al., 2014***).

How, if at all, can group-living animals improve collective decision accuracy while suppressing the potentially deleterious constraint of decision-making biases through trial-and-error learning? One of the strong candidates of explaining this gap is the fact that studies in human social learning in risky decision making have focused only on either the description-based gambles (***Chung et al., 2015***; ***Bault et al., 2011***; ***Suzuki et al., 2016***; ***Shupp and Williams, 2008***) or extreme conformity where individual choices are regulated fully by others’ behaviour (***Denrell and Le Mens, 2007***, ***2016***), but not on experienced-based situations where both individual and social learning affect behavioural outcomes, a form of decision making widespread in group-living animals and humans (***Hertwig and Erev, 2009***; ***Camazine et al., 2001***; ***Toyokawa et al., 2019***).

Here we provide a mathematical model and experimental results of a collective experience-based decision-making task. Using an agent-based computational model simulation and analyses of a differential equation model representing approximated dynamics of collective behaviour, we show theoretically that optimal alternation of behavioural bias thanks to social learning, namely, *collective behavioural rescue*, could emerge through self-organisation because modest conformist social influences can promote information sampling. The theoretical predictions were supported by online behavioural experiments with 585 human subjects. We argue that collective reinforcement learning can alter, rather than amplify, the unfavourable behavioural bias, suggesting a previously overlooked benefit of conformist social influence.

### The Agent-Based Model

We assumed that decision makers could repeatedly choose between a safe alter-native that would provide certain payoffs and a risky alternative that would provide uncertain payoffs (Fig. 1 a), while being able to observe other players’ choices (***McElreath et al., 2005***, ***2008***; ***Toyokawa et al., 2014***, ***2017***, ***2019***; ***Deffner et al., 2020***). To maximise one’s own long-term individual profit in such an experience-based risky decision-making task, it is crucial to strike the right balance between exploiting the option that has seemed better so far and exploring the other options to seek informational gain. Because of the nature of adaptive information sampling under such exploration-exploitation trade-offs, lone decision makers often end up being risk averse, trying to reduce the chance of further failures once the individual has experienced an unfavourable outcome from the risky alternative (***March, 1996***; ***Denrell, 2007***; ***Hertwig and Erev, 2009***), a phenomenon known as the *hot stove effect.* Within the framework of this task, risk aversion is suboptimal in the long run if the risky option provides higher payoffs on average (***Denrell and March, 2001***).

**Figure 1.**
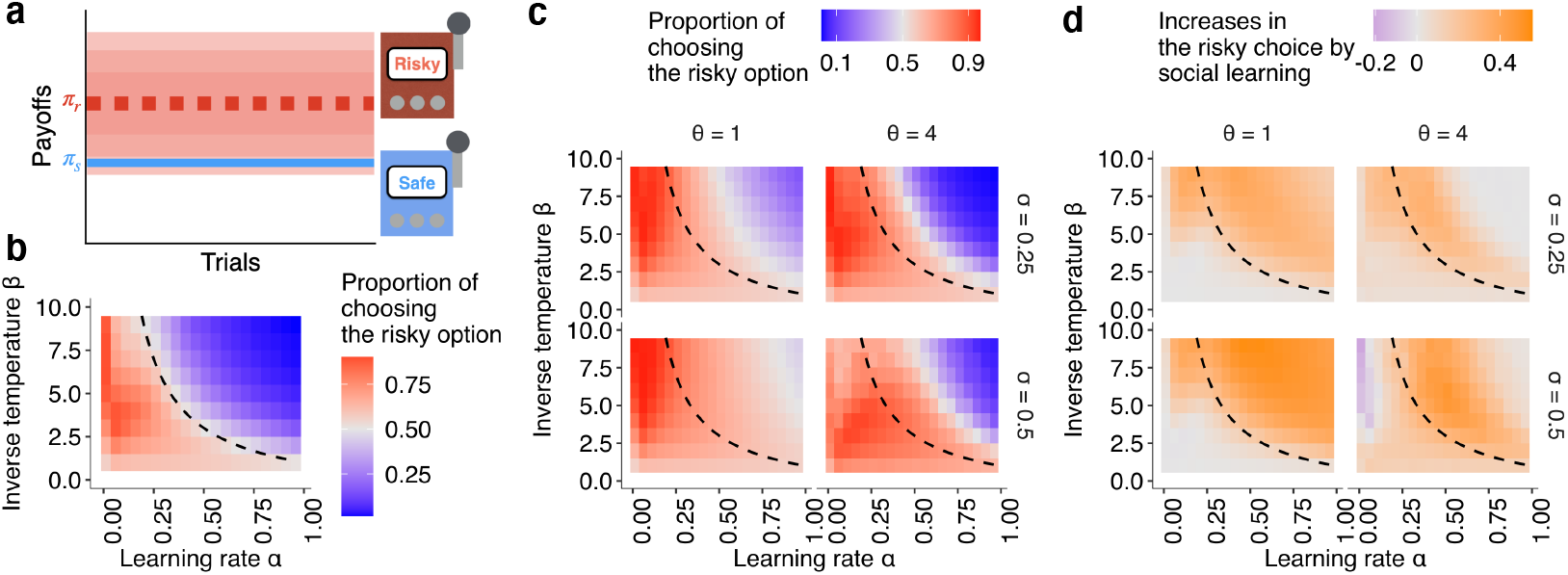
Mitigation of the hot stove effect by collective decision making in a two-armed risky bandit task. (a) A schematic diagram of the task. A safe option provides a constant reward *π_s_* = 1 whereas a risky option provides a reward randomly drawn from a Gaussian distribution with mean *μ* = 1.5 and s.d. = 1. (b, c): The emergence of risk-taking bias due to the hot stove effect depending on a combination of the reinforcement learning parameters;(b): under no social influence (i.e., the copying weight *σ* = 0), and (c): under social influences with different values of the conformity exponents *θ* and copying weights *σ*. The dashed curve is the asymptotic equilibrium at which asocial learners are expected to end up choosing the two alternatives with equal likelihood (i.e., 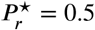), which is given analytically by *β* = (2 – *α*)/*α*. The coloured background is a result of the agent-based simulation with total trials *T* = 150 and group size *N* = 10, showing the average proportion of choosing the risky option in the second half of the learning trials *P*_*r,t*>75_ > 0.5 under a given combination of the parameters. (d): The differences between the mean proportion of risk aversion of asocial learners and that of social learners, highlighting regions in which performance is improved (orange) or undermined (purple) by social learning.

### The baseline asocial learning model and hot stove effect

We considered a sole decision maker facing a repeated choice between a safe (i.e., certain) alternative *s* that generated a constant payoff *π_s_* and a risky (i.e., uncertain) alternative *r* that generated a random payoff *π_r_*, following a Gaussian distribution 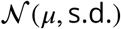 (i.e., a two-armed risky bandit task; Fig. 1a).

We assumed that the decision maker updates their value of choosing the alternative *i* (∈ {*s,r*}) at time *t* following the Rescorla-Wagner learning rule: *Q*_*i,t*+1_ ← (1–*α*)*Q_i,t_*+*απ_i,t_*, where *α* (0 ≤ *α* ≤ 1) is a *learning rate*, manipulating the step size of the belief updating, and *π_i,t_* is a realised payoff from the chosen alternative *i* at time *t* (*Sutton and Barto, 2018).* The larger the *α*, the more weight is given to recent experiences, making reinforcement learning more myopic. The *Q* value for the unchosen alternative is unchanged. Before the first choice, individuals had no previous preference for either option (i.e., *Q*_*r*,1_ = *Q*_*s*,1_ = 0). Then *Q* values were translated into choice probabilities through a softmax (or multinomial-logistic) function such that *P_i,t_* = exp(*βQ_i,t_*)/(exp(*βQ_s,t_*) + exp(*βQ_r,t_*)), where *β*, an *inverse temperature*, is a parameter regulating how sensitive the choice probability is to the value of the estimate *Q* (i.e., controlling the proneness to explore). As *β* → 0, the softmax probability approximates to a random choice (i.e., highly explorative). Conversely, if *β* → +∞, it asymptotes to a deterministic choice in favour of the option with the highest *Q* value (i.e., highly exploitative).

In such a risk-heterogeneous multi-armed bandit setting, reinforcement learners are prone to exhibiting suboptimal risk aversion (***March, 1996***; ***Denrell, 2007***; ***Hertwig and Erev, 2009***), even though they could have achieved high performance in a risk-homogeneous task where all options have an equivalent payoff variance (***Sutton and Barto, 2018***). ***Denrell*** (***2007***) mathematically derived a condition under which suboptimal risk aversion arises, depicted by the dashed curve in Fig. 1b. The suboptimal risk aversion becomes prominent when value updating in learning is myopic (i.e., when the learning rate *α* is large) and/or action selection is exploitative (i.e., when the inverse temperature of the softmax action selection *β* is large). Under such circumstances, the hot stove effect occurs: Experiences of low-value payoffs from the risky option tend to discourage decision makers from further choosing the risky option, trapping them in the safe alternative. In sum, whenever the interaction between the two learning parameters *α*(*β* + 1) exceeds a threshold value, which was 2 in the current example, decision makers are expected to become averse to the risky option (Fig. 2).

**Figure 2.**
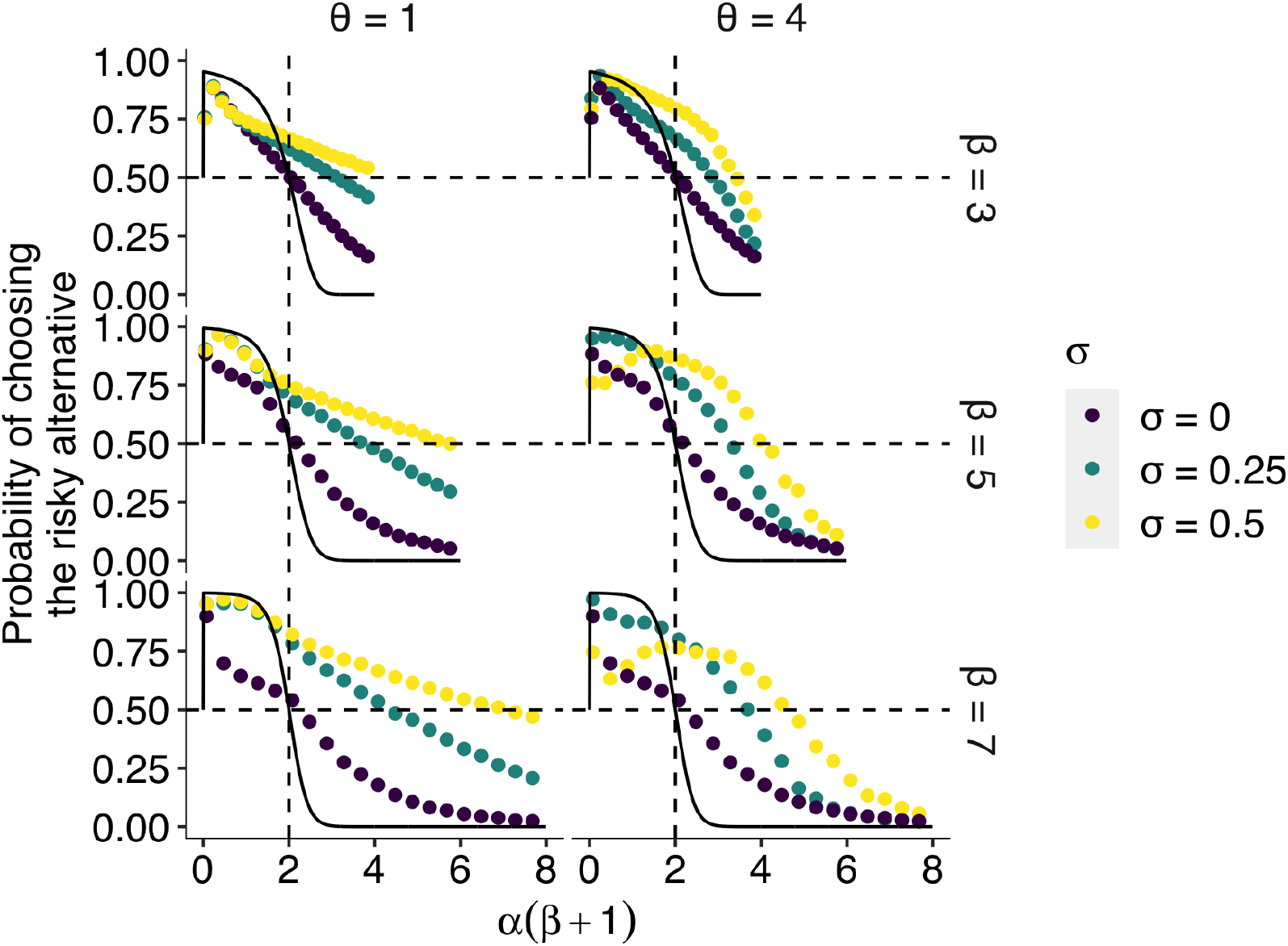
The effect of social learning on average decision performance. The *x* axis is a product of two reinforcement learning parameters *α*(*β* + 1), namely, the susceptibility to the hot stove effect. The *y* axis is the mean probability of choosing the optimal risky alternative in the last 75 trials in a two-armed bandit task whose setup was the same as in Fig. 1. The black solid curve is the analytical prediction of the asymptotic performance of individual reinforcement learning with infinite time horizon *T* → +∞. The analytical curve shows a choice shift emerging at *α*(*β* + 1) = 2; that is, individual learners ultimately prefer the safe to the risky option when *α*(*β* +1) > 2 holds in the current setup of the task. The dotted curves are mean results of agent-based simulations of social learners with two different mean values of the copying weight *σ* ∈ {0.25,0.5} (green and yellow, respectively) and asocial learners with *σ* = 0 (purple). The difference between the agent-based simulation with *σ* = 0 and the analytical result was due to the finite number of decision trials in the simulation, and hence, the longer the horizon, the closer they become (Supplementary Fig. 12). Each panel shows a different combination of the inverse temperature *β* and the conformity exponent *θ.*

In the main analysis, we focused on the case where the risky alternative had *μ* = 1.5 and s.d. = 1 while the safe alternative generates *π_s_* = 1, that is, where choosing the risky alternative was the optimal strategy for a decision maker in the long run.

### Collective learning and social influences

Using the simplest setup of the two-armed bandit (***March, 1996***; ***Denrell, 2007***), we considered a collective learning situation in which a group of 10 individuals complete the task simultaneously and individuals can obtain social information. For social information, we assumed a simple frequency-based social cue specifying distributions of individual choices (***McElreath et al., 2005***, ***2008***; ***Toyokawa et al., 2017***, ***2019***; ***Deffner et al., 2020***). We assumed that individuals could not observe others’ earnings, ensuring that individuals could not learn from approximate forgone payoffs that would have easily rescued people from the hot stove effect (***Denrell, 2007***; ***Yechiam and Busemeyer, 2006***).

A payoff realised was independent of others’ decisions and it was drawn solely from the payoff probability distribution specific to each alternative, thereby we assumed neither direct social competitions over the monetary reward (***Giraldeau and Caraco, 2000***) nor normative pressures towards majority alignment (***Cialdini and Goldstein, 2004***; ***Mahmoodi et al., 2018***). The value of social information was assumed to be only informational (***Nakahashi, 2007***). Nevertheless, our model could apply to the context of normative social influences, because what we assumed here was modifications in individual choice probabilities due to social influences, irrespective of underlying motivations of conformity.

Following the previous modelling of social learning in such multi-agent multi-armed bandit situations (e.g., ***Aplin et al., 2017***; ***Barrett et al., 2017***; ***McElreath et al., 2005***, ***2008***; ***Toyokawa et al., 2017***, ***2019***; ***Deffner et al., 2020***), we assumed that social influences on reinforcement learning would be expressed as a weighted average between the softmax probability based on the *Q* values and the conformist social influence as follows:

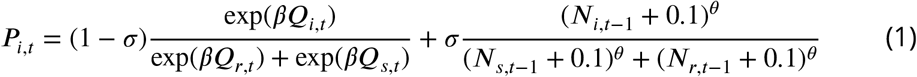

where *σ* was a weight given to the social influence (*copying weight*) and *θ* was the strength of conformist influence (*conformity exponent*), which determines the influence of social frequency of choosing the alternative *i* at time *t*–1, that is, *N*_*i,t*–1_. The larger the conformity exponent *θ*, the higher the influence that was given to an alternative that was chosen by more individuals, with non-linear conformist social influence arising when *θ* > 1. Note that there was no actual social influence when *θ* = 0 because in this case the ‘social influence’ favours a uniformly random choice, irrespective of the social frequency distribution *N*_s_ and *N_r_*. We added a small number, 0.1, to *N*_*i,t*–1_ so that an option chosen by no one (i.e., *N*_*i,t*–1_ = 0) could provide the highest social influence when *θ* < 0 (negative frequency bias). Although this additional 0.1 slightly reduces the conformity influence when *θ* > 0, we confirmed that the results were qualitatively unchanged. Note also that in the first trial *t* = 1, we assumed that the choice was determined solely by the asocial softmax function because there was no social information available yet.

Note that, when *σ* = 0, there is no social influence, and the decision maker is considered as an asocial learner. It is also worth noting that, when *σ* =1 with *θ* > 0, individual choices are assumed to be contingent fully upon majority’s behaviour, which was assumed in some previous models of strong conformist social influences in sampling behaviour (***Denrell andLe Mens, 2016***). Our model is a natural extension of both the asocial reinforcement learning and the model of ‘extreme conformity’, as these conditions can be expressed as a special case of parameter combinations. We will discuss the implications of this extension in the Discussion. The descriptions of the parameters are shown in Table 1.

**Table 1.** Summary of the learning model parameters

| Symbol | Meaning | Range of the value |
| --- | --- | --- |
| $\alpha$ | Learning rate | [0, 1] |
| $\beta$ | Inverse temperature | [0, $+\infty$] |
| $\alpha(1 + \beta)$ | Susceptibility to the hot stove effect | |
| $\sigma$ | Copying weight | [0, 1] |
| $\theta$ | Conformity exponent | $[-\infty, +\infty]$ |

## Results

### The collective behavioural rescue effect

We observed a mitigation of suboptimal risk aversion by social influences in the simplest two-armed bandit task. As shown in Fig. 1c, a simple frequency-based social influence, even with a conformist influence (*θ* > 1), widened the region of parameter combinations where decision makers could escape from the suboptimal risk aversion (the red area). The increment of the area of adaptive risk seeking was greater with *θ* = 1 than *θ* = 4. When *θ* = 1, a large copying weight *σ* can eliminate almost all the area of risk aversion (Fig. 1 c; see also Supplementary Fig. 7 for a greater range of parameter combinations). Note that increasing the copying weight *σ* → 1 eventually approximates a random choice (Supplementary Fig. 7). Interestingly, such a switch to risk seeking did not emerge when risk aversion was actually optimal (Supplementary Fig. 9), suggesting that social influence does not always increase risk seeking;instead, the effect seems to be more prominent especially when risk seeking is beneficial in the long run.

Fig. 2 highlights the extent to which risk aversion was relaxed through social influences. Individuals with positive *σ* > 0 could maintain a high proportion of risk seeking even in the region of high susceptibility to the hot stove effect *α*(*β* + 1) > 2. Although social learners eventually fell into a risk-averse regime with increasing *α*(*β* + 1), risk aversion was largely mitigated compared to the performance of individual learners who had *σ* = 0. Interestingly, the probability of choosing the optimal risky option was maximised at an intermediate value of *α*(*β* + 1) when the conformity exponent was large *θ* = 4 and the copying weight was high *σ* = 0.5.

In the region of less susceptibility to the hot stove effect (*α*(*β* + 1) < 2), social influence could enhance individual optimal risk seeking up to the theoretical benchmark expected in individual reinforcement learning with an infinite time horizon. A socially induced increase in risk seeking in the region *α*(*β* +1) < 2 was more evident with larger *β*, and hence with smaller *α* to satisfy *α*(*β* +1) < 2. The smaller the learning rate *α*, the longer it would take to achieve the asymptotic equilibrium state, due to slow value updating. Asocial learners, as well as social learners with high *σ* (= 0.5) coupled with high *θ* (= 4), were still far from the analytical benchmark, whereas social learners with weak social influence *σ* = 0.25 were nearly able to converge on the benchmark performance, suggesting that social learning might affect the speed of learning. Indeed, a longer time horizon *T* = 1075 reduced the advantage of weak social learners in this *α*(*β* +1) < 2 region because slow learners could now achieve the benchmark accuracy (Supplementary Figs. 12 and 13).

Approaching the benchmark with an elongated time horizon, and the con-comitant reduction in the advantage of social learners, was also found in the high susceptibility region *α*(*β* + 1) ≫ 2 especially for those who had a high conformity exponent *θ* = 4 (Supplementary Fig. 12). Notably, however, facilitation of optimal risk seeking became further evident in the other intermediate region 2 < *α*(*β* + 1) < 4. This suggests that merely speeding up or slowing down learning could not satisfactorily account for the qualitative choice shift emerging through social influences.

We obtained similar results across different settings of the multi-armed ban-dit task, such as a skewed payoff distribution in which either large or small payoffs were randomly drawn from a Bernoulli process (***March, 1996***; ***Denrell, 2007***) (Supplementary Figs. 10) and increased option numbers (Supplementary Figs. 11). Further, the conclusion still held under different assumptions of social influences on reinforcement learning, assuming that the conformist influence acts on the learning process (the value-shaping model (***Najar et al., 2020***)) rather than on the action-selection process (the decision-biasing model) assumed above (see Supplementary Methods). Although groups of agents with the value-shaping algorithm seemed more prone to herding, the results were not qualitatively changed and collective behavioural rescue emerged when *β* was sufficiently small (Supplementary Figs. 8).

### The effect of individual heterogeneity

We have thus far assumed no parameter variations across individuals in a group to focus on the qualitative differences between social and asocial learners’ behaviour. However, individual differences in development, state, or experience or variations in behaviour caused by personality traits might either facilitate or undermine collective decision performance. Especially if a group is composed of both types of individuals, those who are less susceptible to the hot stove effect *α*(*β* + 1) < 2 as well as those who are more susceptible *α*(*β* + 1) > 2, it remains unclear who benefits from the rescue effect: Is it only those individuals with *α*(*β* + 1) > 2 who enjoy the benefit, or can collective intelligence benefit a group as a whole? For the sake of simplicity, we considered groups of five individuals here.

Fig. 3a shows the effect of heterogeneity in the learning rate *α*. Heterogeneous groups (shown in green, blue, or purple) performed better on average than a homogeneous group (represented by the yellow diamond). The heterogeneous groups owed this overall improvement to the large rescue effect operating for highly susceptible individuals who had *α*(*β* + 1) ≫ 2. On the other hand, the performance of less susceptible individuals [*α*(*β* + 1) < 2] was slightly undermined compared to the asocial benchmark performance. Notably, however, how large the detrimental effect was for the low-susceptibility individuals depended on the group’s composition: The undermining effect was largely mitigated when low-susceptibility individuals [*α*(*β* + 1) < 2] made up a majority of a group (3 of 5; the blue line), whereas they performed worse than the asocial benchmark when the majority were those with high susceptibility (purple).

**Figure 3.**
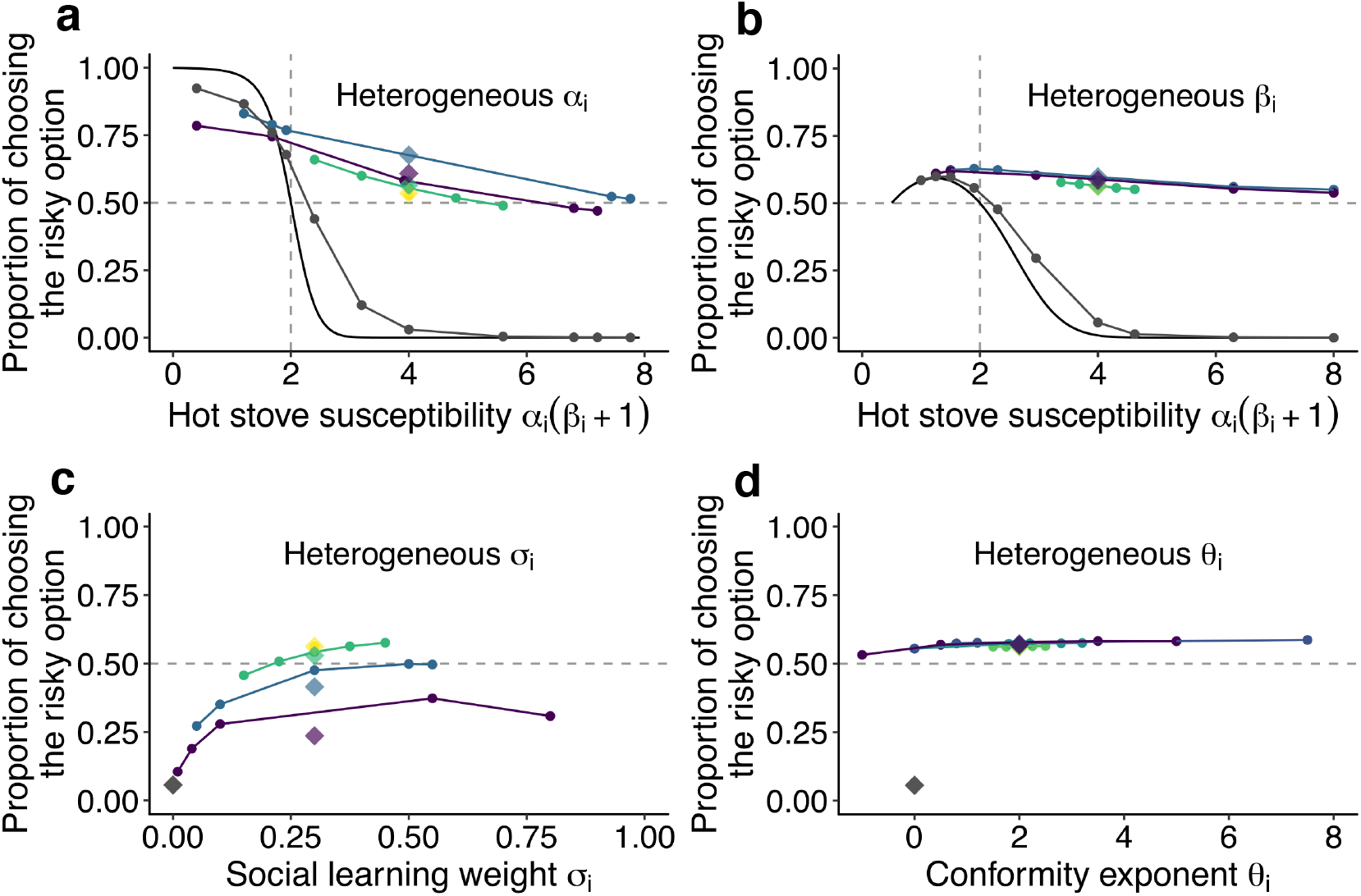
The effect of individual heterogeneity on the proportion of choosing the risky option in the two-armed bandit task. (a) The effect of heterogeneity of *α*, (b) *β*, (c) *σ*, and (d) *θ*. Individual values of a focal behavioural parameter were varied across individuals in a group of five. Other non-focal parameters were identical across individuals within a group. The basic parameter values assigned to non-focal parameters were *α* = 0.5, *β* = 7, *σ* = 0.3, and *θ* = 2, and groups’ mean values of the various focal parameters were matched to these basic values. We simulated 3 different heterogeneous compositions: The majority (3 of 5 individuals) potentially suffered the hot stove effect *α_i_*(*β_i_* + 1) > 2 (a, b) or had the highest diversity in social learning parameters (c, d;purple);the majority were able to overcome the hot stove effect *α_i_*(*β_i_* + 1) < 2 (a, b) or had moderate heterogeneity in the social learning parameters (c, d;blue);and all individuals had *α_i_*(*β_i_* + 1) > 2 but smaller heterogeneity (green). The yellow diamond shows the homogeneous groups’ performance. Lines are drawn through average results across the same compositional groups. Each round dot represents a group member’s mean performance. The diamonds are the average performance of each group. For comparison, asocial learners’ performance, with which the performance of social learners can be evaluated, is shown in grey. For heterogeneous *α* and *β*, the analytical solution of asocial learning performance is shown as a solid-line curve. We ran 20,000 replications for each group composition.

A similar pattern was found for the inverse temperature *β*, although the impact of its heterogeneity was much smaller than that in *α*, and there was little difference between homogeneous and heterogeneous groups (Fig. 3b). Interestingly, no detrimental effect for less susceptible individuals was found in association with the *β* variations.

On the other hand, individual variations in the copying weight *σ* had an overall detrimental effect on collective performance, although individuals in the highest diversity group could still perform better than the asocial benchmark (Fig. 3c). Individuals who had an intermediate level of copying weight achieved relatively higher performance within the group than those who had either higher or lower weight. This was because individuals with lower *σ* could benefit less from social information, while those with higher *σ* relied so heavily on social frequency information that behaviour was barely informed by individual learning, resulting in maladaptive herding or collective illusion (***Denrell and Le Mens, 2016***; ***Toyokawa et al., 2019***). As a result, the average performance decreased with increasing diversity in *σ*.

Such a substantial effect of individual differences was not observed in the conformity exponent *θ* (Fig. 3d), where individual performance was almost stable regardless of whether the individual was heavily conformist (*θ_i_* = 8) or even anticonformist (*θ_i_* = −1). The existence of a few conformists in a group could not itself trigger positive feedback among the group unless other individuals also relied on social information in a conformist-biased way, because the flexible behaviour of non-conformists could keep the group’s distribution nearly flat (i.e., *N_s_* ≈ *N_r_*). Therefore, the existence of individuals with small *θ* in a heterogeneous group could prevent the strong positive feedback from being immediately elicited, compensating for the potential detrimental effect of maladaptive herding by strong conformists.

In sum, the relaxation of, and possibly the complete rescue from, a suboptimal bias in repeated risky decision making emerged in a range of conditions in collective learning. It was not likely a mere speeding up or slowing down of learning process (Supplementary Figs. 12 and 13), nor just an averaging process mixing performances of both risk seekers and risk-averse individuals (Fig. 3). It depended neither on specific characteristics of social learning models (Supplementary Fig. 8) nor on the profile of the bandit task’s setups (Supplementary Fig. 10). Instead, our simulation suggests that self-organisation may play a key role in this emergent phenomenon. To seek a general mechanism underlyingthe observed collective behavioural rescue, in the next section we show a reduced, approximated differential equation model that can provide qualitative insights into the collective decision-making dynamics observed above.

### The simplified collective dynamics model

To obtain a qualitative understanding of self-organisation that seems responsible for the pattern of adaptive behavioural shift observed in our individual-based simulation, we made a reduced model that approximates temporal changes of behaviour of an ‘average’ individual, where the computational details of reinforcement learning were purposely ignored. Specifically, we considered a differential equation model that focuses only on increases and decreases in the expected number of individuals who chose the risky option and the safe option (Fig. 4a). Such a dynamic modelling approach has been commonly used in population ecology and collective animal behaviour research and has proven highly useful in disentangling the factors underlying complex systems (e.g., ***Beckers et al., 1990***; ***Goss et al., 1989***; ***Seeley et al., 1991***; ***Sumpter and Pratt, 2003***; ***Camazine et al., 2001***). The full details of this dynamics model are shown in the Method and Table 3.

To confirm that this approximated model can successfully replicate the funda-mental property of the hot stove effect, we first describe the asocial behavioural model without social influence (see Methods). The baseline, asocial dynamic system has a locally stable non-trivial equilibrium that gives 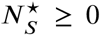 and 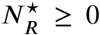, where *N*^★^ means the equilibrium density at which the system stops changing 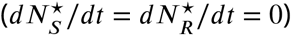. At equilibrium, the ratio between the number of individuals choosing the safe option *S* and the number choosing the risky option *R* is given by 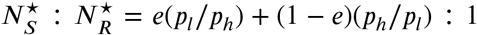, indicating that risk aversion (defined as the case where more people choose the safe option; 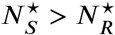) emerges when the inequality 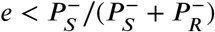 holds.

Fig. 4b visually shows that the population is indeed attracted to the safe option *S*, which results in 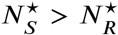, in a wide range of the parameter region even when there is a positive ‘risk premium’ defined as *e* > 1/2. Although individuals choosing the risky option *R* are more likely to become enchanted with the risky option (i.e., choosing *R* makes the belief state positive +) than to get disappointed (i.e., choosing *R* makes the belief state negative –), the risk-seeking equilibrium (defined as 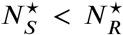) becomes less likely to emerge as the exploration rate *p_l_* decreases, consistent with the hot stove effect caused by asymmetric adaptive sampling (***Denrell, 2007***). Risk seeking never emerges when *e* ≤ 1/2, which is also consistent with the results of reinforcement learning.

This dynamics model provides an illustrative understanding of how the asymmetry of adaptive sampling causes the hot stove effect. Consider the case of high inequality between exploitation (*p_h_*) and exploration (*p_l_*), namely, *p_h_* ≫ *p_l_*. Under such a condition, the behavioural state *S*^−^, that is choosing the safe option with the negative inner belief −, becomes a ‘dead end’ from which individuals can seldom escape once entered. However, if the inequality *p_h_* ≥ *p_l_* is not so large that a substantial fraction of the population now comes back to *R*^−^ from *S*^−^, the increasing number of people belonging to *R*^+^ (that is, 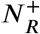) could eventually exceed the number of people ‘spilling out’ to *S*^−^. Such an illustrative analysis shows that the hot stove effect can be overcome if the number of people who get stuck in the dead end *S*^−^ can somehow be reduced. And this is possible if one can increase the ‘come-back’ to *R*^−^. In other words, if any mechanisms can increase 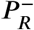 in relation to 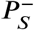, the hot stove effect should be overcome.

Through numerical analyses of the dynamics model with social influence (see Methods), we have confirmed that social influence can indeed increase the flowback rate 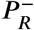, which raises the possibility of risk-seeking equilibrium 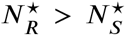 (Fig. 4d; see Supplementary Fig. 14 for a wider parameter region). For an approximation of the bifurcation analysis, we recorded the equilibrium density of the risky state 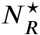 starting from various initial population distributions (that is, varying *N*_*R,t*=0_ and *N*_*S,t*=0_). Fig. 5 shows the conditions under which the system ends up in risk-seeking equilibrium. When the conformity exponent *θ* is not too large (*θ* < 10), risk seeking can be a unique equilibrium irrespective of the initial distribution, attracting the population even from an extremely biased initial distribution such as *N*_*R,t*=0_ = 0 (Fig. 5).

**Figure 4.**
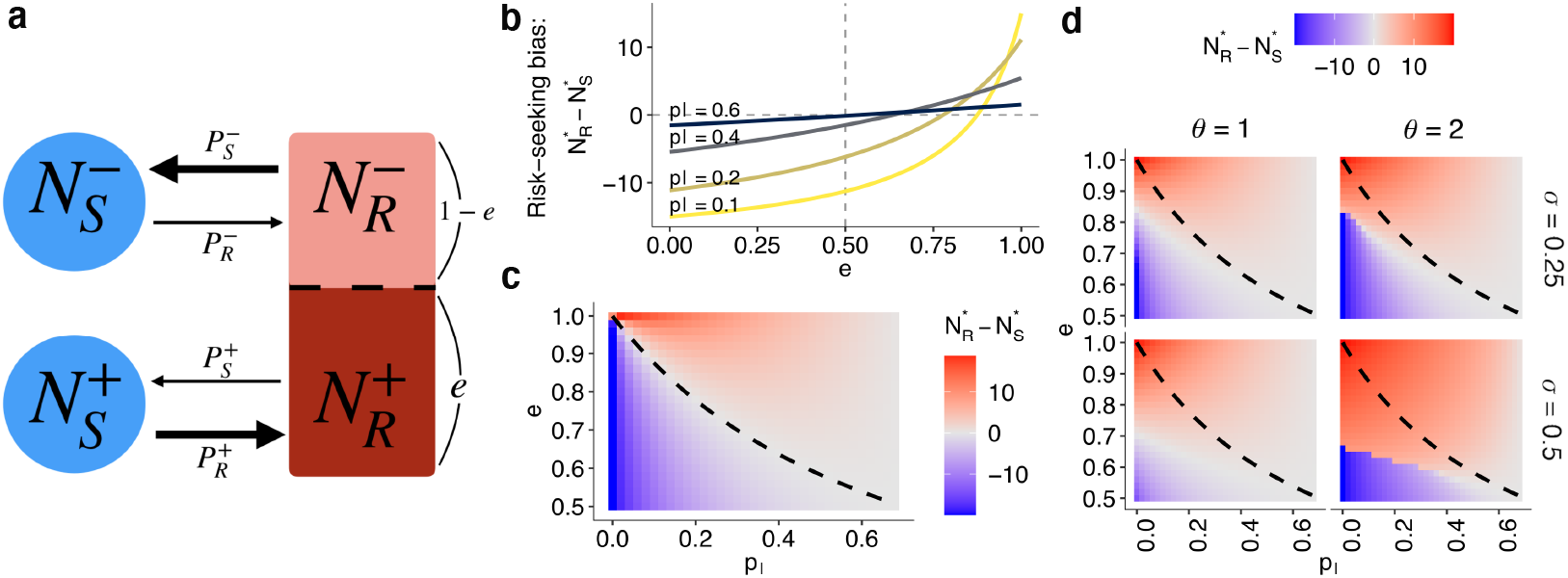
The collective behavioural dynamics model. (a) A schematic diagram of the dynamics. Solid arrows represent a change in population density between connected states at a time step. The thicker the arrow, the larger the per-capita rate of behavioural change. (b, c) The results of the asocial, baseline model where 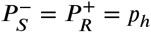 and 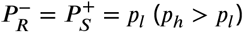. Both figures show the equilibrium bias towards risk seeking (i.e., 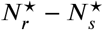) as a function of the degree of risk premium *e* as well as of the per-capita probability of moving to the less preferred behavioural option *p*_1_. (b) The explicit form of the curve is given by –*n*(*p_h_* – *p_l_*) {(1 – *e*)*p_h_* – *e_pl_* } /(*p_h_* + *p_l_*) {1 – *e*)*p_h_* + *e_pl_*}. (c) The dashed curve is the analytically derived neutral equilibrium of the asocial system that results in 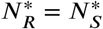, given by *e* = *p_h_*/(*p_h_* + *p_l_*). (d) The equilibrium of the collective behavioural dynamics with social influences. The numerical results were obtained with 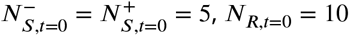, and *p_h_* = 0.7.

Under the conformist bias *θ* ≥ 2, two locally stable equilibria exist. Strong positive feedback dominates the system when both *σ* and *θ* are large. Therefore, the system can end up in either of the equilibria depending solely on the initial density distribution, consistent with the conventional view of herding (***Denrell and Le Mens, 2016***; ***Toyokawa et al., 2019***). This is also consistent with a well-known result of collective foraging by pheromone trail ants, which react to social information is a conformity-like manner (***Beckers et al., 1990***; ***Camazine et al., 2001***).

Notably, however, even with a large conformist bias, such as *θ* = 2, there is a region with a moderate value of *σ* where risk seeking remains a unique equilibrium when the premium of the risky option *e* was high (*e* ≥ 0.7). In this regime, the benefit of collective behavioural rescue can dominate without any possibility of maladaptive herding.

It is worth noting that in the case of *θ* = 0, where individuals make merely a random choice at a rate *σ*, risk aversion is also relaxed (Fig. 5, the leftmost column), and the adaptive risky shift even emerges around 0.25 < *σ* < 1. However, this ostensible behavioural rescue is due solely to the pure effect of additional random exploration that reduces 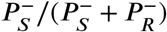, mitigating stickiness to the dead-end status *S*^−^. When *σ* → 1 with *θ* = 0, therefore, the risky shift eventually disappears because the individuals choose between *S* and *R* almost randomly.

However, the collective risky shift observed in the conditions of *θ* > 0 cannot be explained solely by the mere addition of exploration. A weak conformist bias (i.e., a linear response to the social frequency; *θ* = 1) monotonically increases the equilibrium density 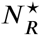 with increasing social influence *σ*, which goes beyond the level of risky shift observed with the addition of random choice (Fig. 5). Therefore, although the collective rescue might indeed owe its part of the mitigation of the hot stove effect to increasing exploration, the further enhancement of risk seeking cannot be fully explained by it alone.

The key is the interaction between negative and positive feedback. As we discussed above, risk aversion is reduced if the ratio 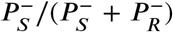 decreases, either by increasing 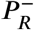 or reducing 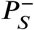. The per individual probability of choosing the safe option with belief −, that is, 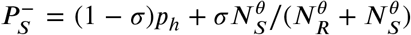, becomes smaller than the baseline exploitation probability *p_h_*, when 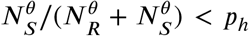. Even though the majority of the population may still choose the safe alternative and hence *N_S_* > *N_R_*, the inequality 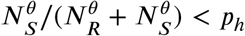 can nevertheless hold if one takes a sufficiently small value of *θ*. Crucially, the reduction of 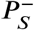 leads to a further reduction of 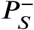 itself through decreasing 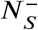, thereby further decreasing the social influence supporting the safe option 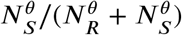. Such a negative feedback process weakens the concomitant risk aversion. Naturally, this negative feedback can be maximised with *θ* = 0.

Once the negative feedback has weakened the underlying risk aversion, the majority of the population may eventually choose the risky option, an effect evident in the case of *θ* = 0 (Fig. 5). What uniquely operates in cases of *θ* > 0 is that because *N_S_* is a minority by now, positive feedback starts. Thanks to the conformist bias, the inequality *N_R_* > *N_S_* is further *amplified.* In this phase, the larger *θ*, the stronger the concomitant relationship 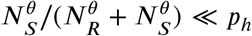. Such positive feedback will never operate with *θ* ≤ 0.

In conclusion, it is the synergy of negative and positive feedback that explains the full range of adaptive risky shift. Neither positive nor negative feedback alone can account for both accuracy and flexibility emerging through collective learning and decision making. The results are qualitatively unchanged across a range of different combinations of *e*, *p_l_*, and *p_h_* (Supporting Figs. 14 and 15). It is worth noting that when *e* < 0.5, the system tends to exhibit risk aversion (Supplementary Fig. 15), consistent with the result of the agent-based simulation for the case where the mean payoff of the risky option was smaller than that of the safe option (Supplementary Fig. 9). Therefore, the system does not mindlessly prefer risk seeking, but it becomes risk prone only when to do so is favourable in the long run.

### An experimental demonstration

To investigate whether the collective rescue effect can operate in reality, we con-ducted a series of online behavioural experiments using human participants. The experimental task was basically a replication of the agent-based model described above, although the parameters of the bandit tasks were different (see the Method for the details of the experimental procedures; Supplementary Figure 11). One hundred eighty-five adult human subjects performed the individual task without social interactions, while 400 subjects performed the task collectively with group sizes ranging from 2 to 8 (Supplementary Fig. 17 and 19). We used four different settings for the multi-armed bandit tasks. Three of them were positive risk premium (PRP) tasks that had an optimal risky alternative, while the other was a negative risk premium (NRP) task that had a suboptimal risky alternative (see Methods).

The behavioural results with statistical model fitting confirmed the predictions of the theoretical model. In the PRP task subjects who had a larger estimated value of the susceptibility to the hot stove effect (*α_i_*(*β_i_* + 1)) were less likely to choose the risky alternative, whereas those who had a smaller value of *α_i_*(*β_i_* + 1) had a higher chance of choosing the safe alternative (Fig. 6a-c), consistent with the theory of the hot stove effect (Fig. 2, Supplementary Fig. 11). In the NRP task, individuals tended to choose the favourable safe option more often than they chose the risky option in a range of the susceptibility value *α_i_*(*β_i_*, + 1) (Fig. 6d), which was also consistent with the model prediction (Supplementary Fig. 18). These results suggest that our model captured subjects’ behaviour well. We confirmed a high performance of parameter recovery in our model fitting procedure (Supplementary Fig. 16).

**Figure 5.**
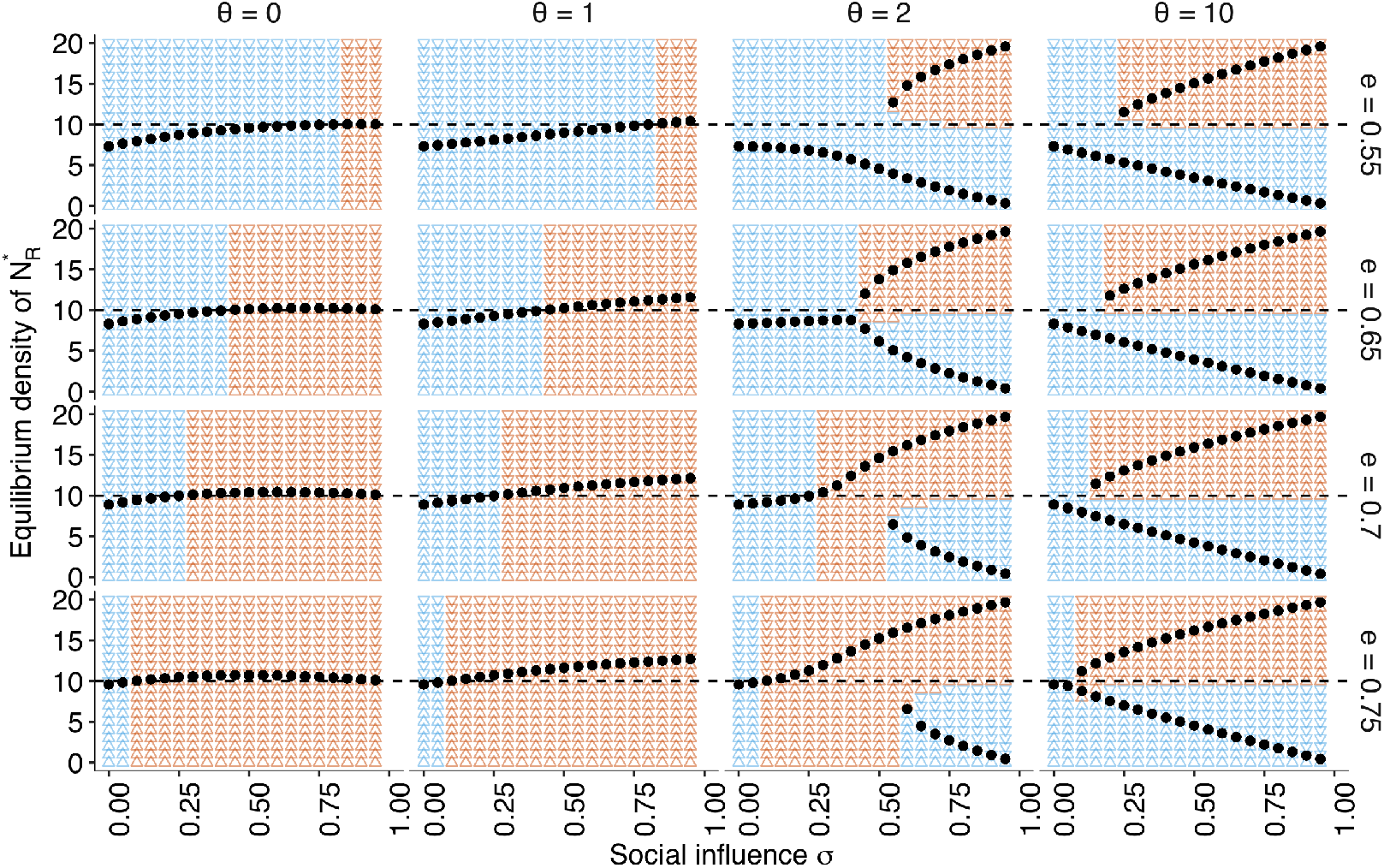
The approximate bifurcation analysis. The relationships between the social influence weight *σ* and the equilibrium number of individuals in the risky behavioural state 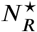 across different conformity exponents *θ* ∈ {0, 1, 2, 10} and different values of risk premium *e* ∈ {0.55,0.65,0.7,0.75}, are shown as black dots. The background colours indicate regions where the system approaches either risk aversion (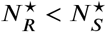; blue) or risk seeking (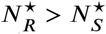; red). The horizontal dashed line is *N_R_* = *N_S_* = 10. Two locally stable equilibria emerge when *θ* = 2 and *θ* = 10, which suggests that the system has a bifurcation when *σ* is sufficiently large. The other parameters are set to *p_h_* = 0.7, *p_l_* = 0.2, and *N* = 20.

**Figure 6.**
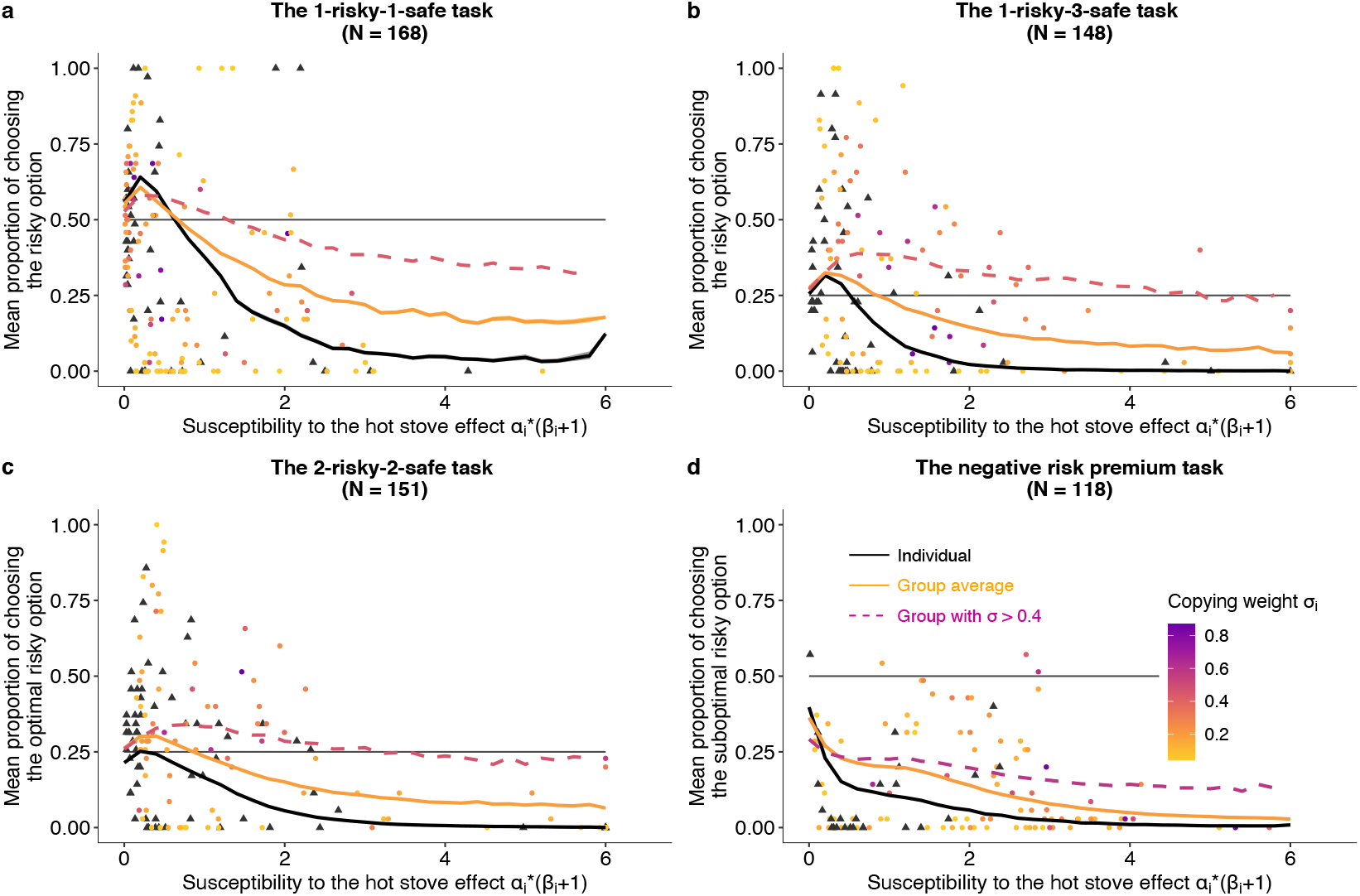
The effect of social learning on decision performance in the online behavioural experiments. (a) A two-armed bandit task (*N* = 168). (b) A 1-risky-3-safe (four-armed) bandit task (*N* = 148). (c) A 2-risky-2-safe (four-armed) bandit task (*N* = 151). (d) A negative risk premium two-armed bandit task (*N* = 118). The *x* axis is *α*,(*β_i_*, + 1), namely, the susceptibility to the hot stove effect. (a,b,and d) The *y* axis is the mean proportion of choosing the risky alternative averaged over the second half of the trials. (c)*y* axis is the mean proportion of choosing the optimal risky alternative averaged over the second half of the trials. The black triangles represent subjects in the individual learning condition, and the coloured dots are those in the group condition with group sizes ranging from 2 to 8. For subjects in the group condition, the fit value of copying weight *σ_i_* is shown in the colour gradient. The solid lines are predictions from simulations of the model with the calibrated global parameters for the individual condition (black) and for the group condition (orange). The dashed violet curves are hypothetical predictions of the calibrated model supplied with a larger copying weight than that for real human subjects, that is, 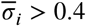.

The behaviour in the group condition supports our theoretical predictions. In the PRP tasks, the proportion of choosing the favourable risky option increased with social influence (*σ_i_*) particularly for individuals who had a high susceptibility to the hot stove effect. On the other hand, social influence had little benefit for those who had a low susceptibility to the hot stove effect (e.g., *α_i_*(*β_i_* + 1) ≤ 0.5).

However, a complete switch of the majority’s behaviour from the suboptimal options to the optimal risky option (i.e., *P_r_* > 0.5 for the two-armed task and *P_r_* > 0.25 for the four-armed task) was not widely observed (Supplementary Fig. 17). This was because of the low reliance on social learning (i.e., the mean *σ_i_*, = 0.19 and the median *σ_i_* = 0.12), coupled with the low susceptibility *α_i_*(*β_i_*+1) in individual learners (i.e., the mean *α_i_*(*β_i_*+1) = 0.8 and median = 0.3 for the individual condition, whereas the mean = 1.1 and median = 0.5 for the group condition). Although the mitigation of risk aversion still emerged for those with higher *α_i_*(*β_i_* + 1), the weak reliance on social learning hindered the strong collective rescue effect because strong positive feedback was not robustly formed.

**Figure 17.**
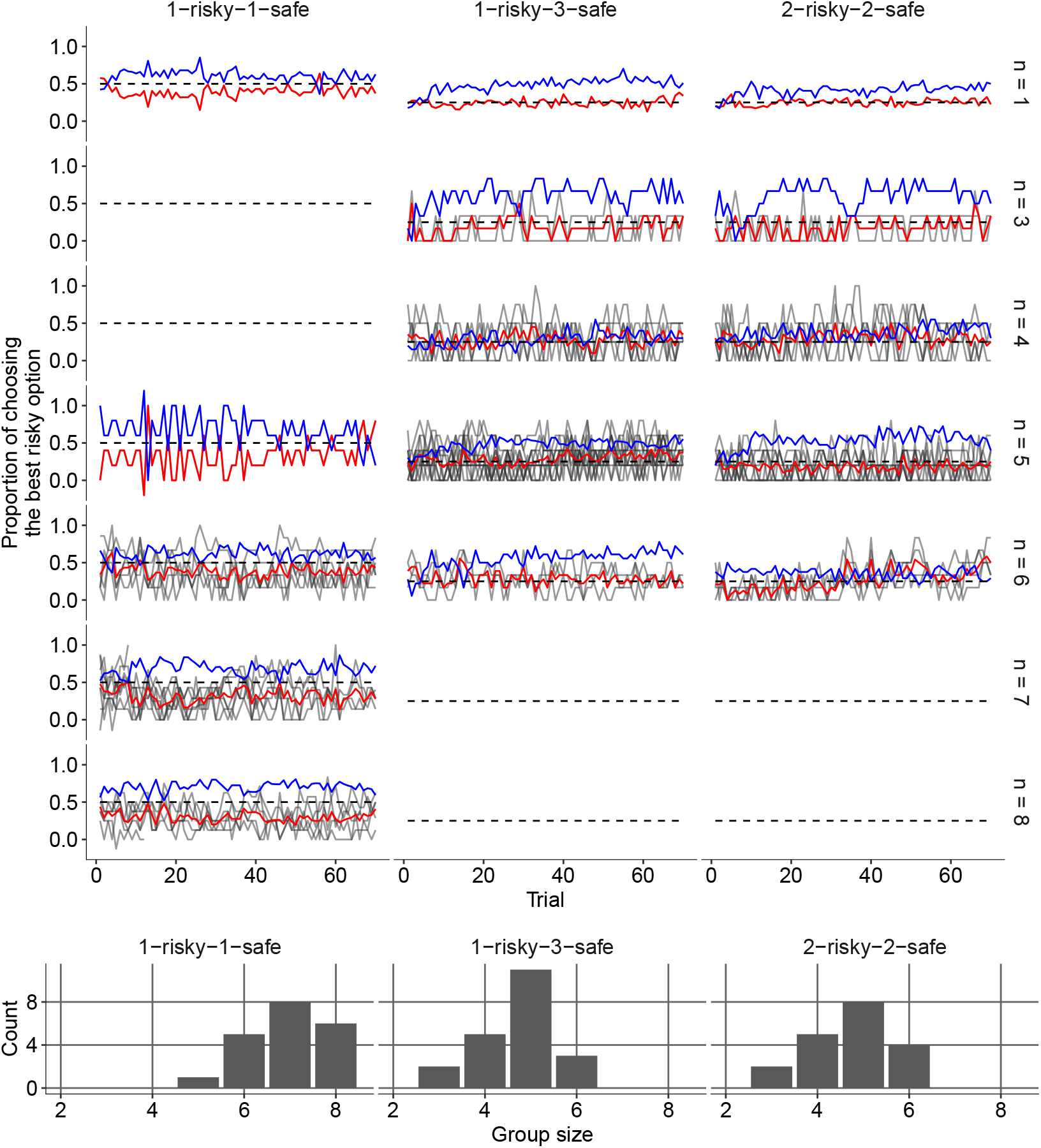
Dynamics of the experimental groups’ mean performance and the distribution of group size. Group size is indicated on the righthand side. Each group’s mean proportion of choosing the best risky option is shown in grey;mean value across the same group-size category is shown in red. The blue line is the mean proportion of choosing the best safe option (that is suboptimal) averaged across the same group-size category.

The estimated global mean of the conformity exponent *θ* was larger than 1 for all tasks (Table 2), suggesting that human groups might still have possessed the potential to set up the positive-feedback loops required for a strong rescue effect. In keeping with this, if we extrapolated the larger value of the copying weight (i.e., 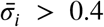) into the best fitting social learning model with the other parameters calibrated, a strong collective rescue became prominent (Fig. 6, the violet dashed lines).

**Table 2.**
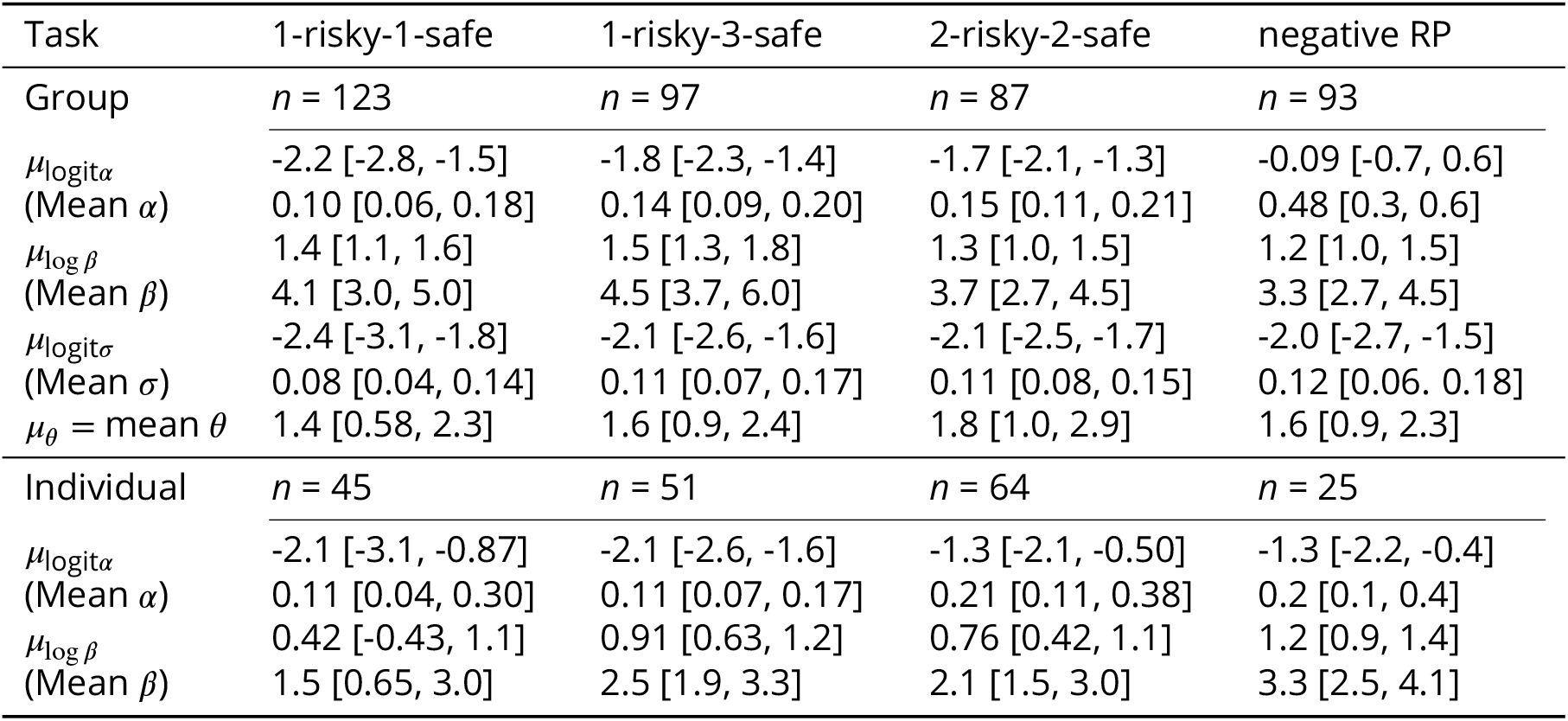
Means and 95% Bayesian credible intervals (shown in square brackets) of the global parameters of the learning model. The group condition and individual condition are shown separately. All parameters satisfied the Gelman-Rubin criterion *R* < 1.01. All estimates are based on over 500 effective samples from the posterior.

**Table 3.**
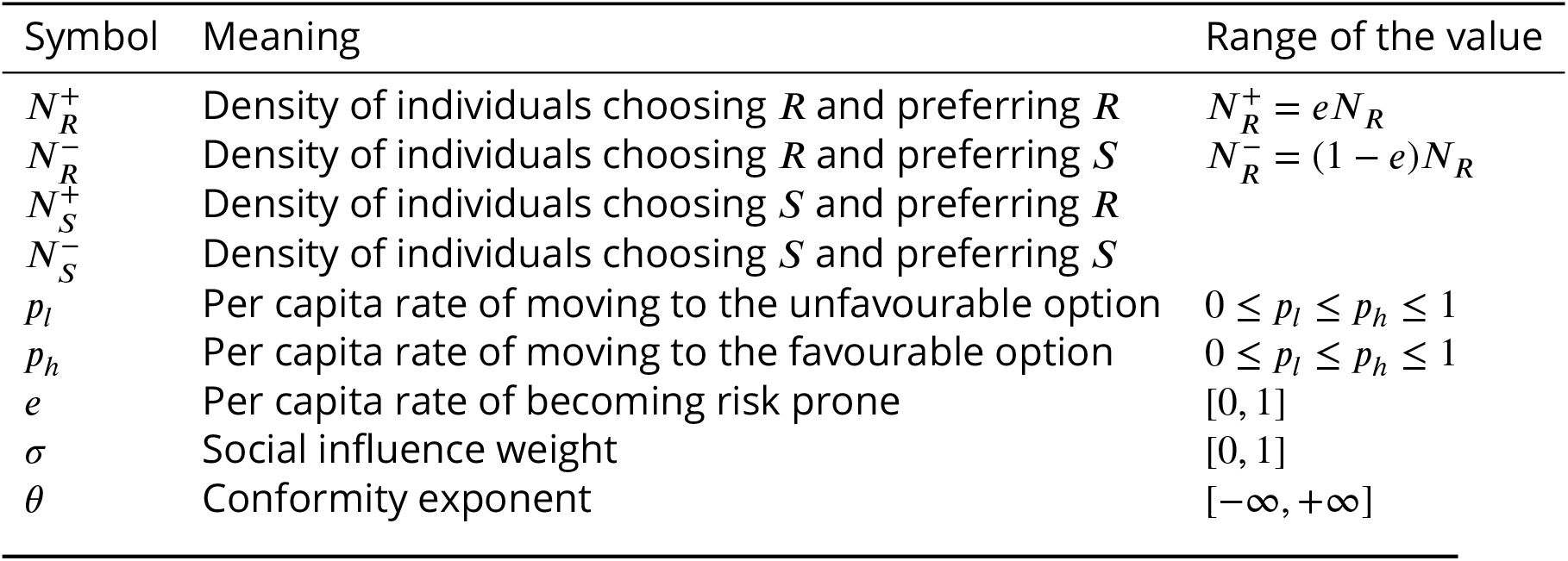
Summary of the differential equation model parameters

In the NRP task, conformist social influence undermined the proportion of choosing the optimal safe option and increased adverse risk seeking, although a complete switch of the majority’s behaviour to the suboptimal risky option did not happen (Fig. 6d; Suppelementary Fig. 19). Such promotion of suboptimal risk taking was particularly prominent when the susceptibility value *α_i_*(*β_i_*+1) was large. Nonetheless, the extent to which risk taking was increased in the NRP condition was smaller than that in the PRP tasks, consistent with our model prediction that conformist social learning is more likely to promote favourable risk taking (Fig. 18). It is worth noting that the estimated learning rates (i.e., mean *α_i_* = 0.48) in the NRP task were larger than that in other PRP tasks (mean *α_i_* < 0.21; Table 2), making social learning particularly deleterious when risk taking is suboptimal (Supplementary Fig. 18). In the discussion, we will discuss about the effect of experimental setting on the human learning strategies, which can be explored in the future studies.

**Figure 18. (Figure 6 - figure supplement 2).**
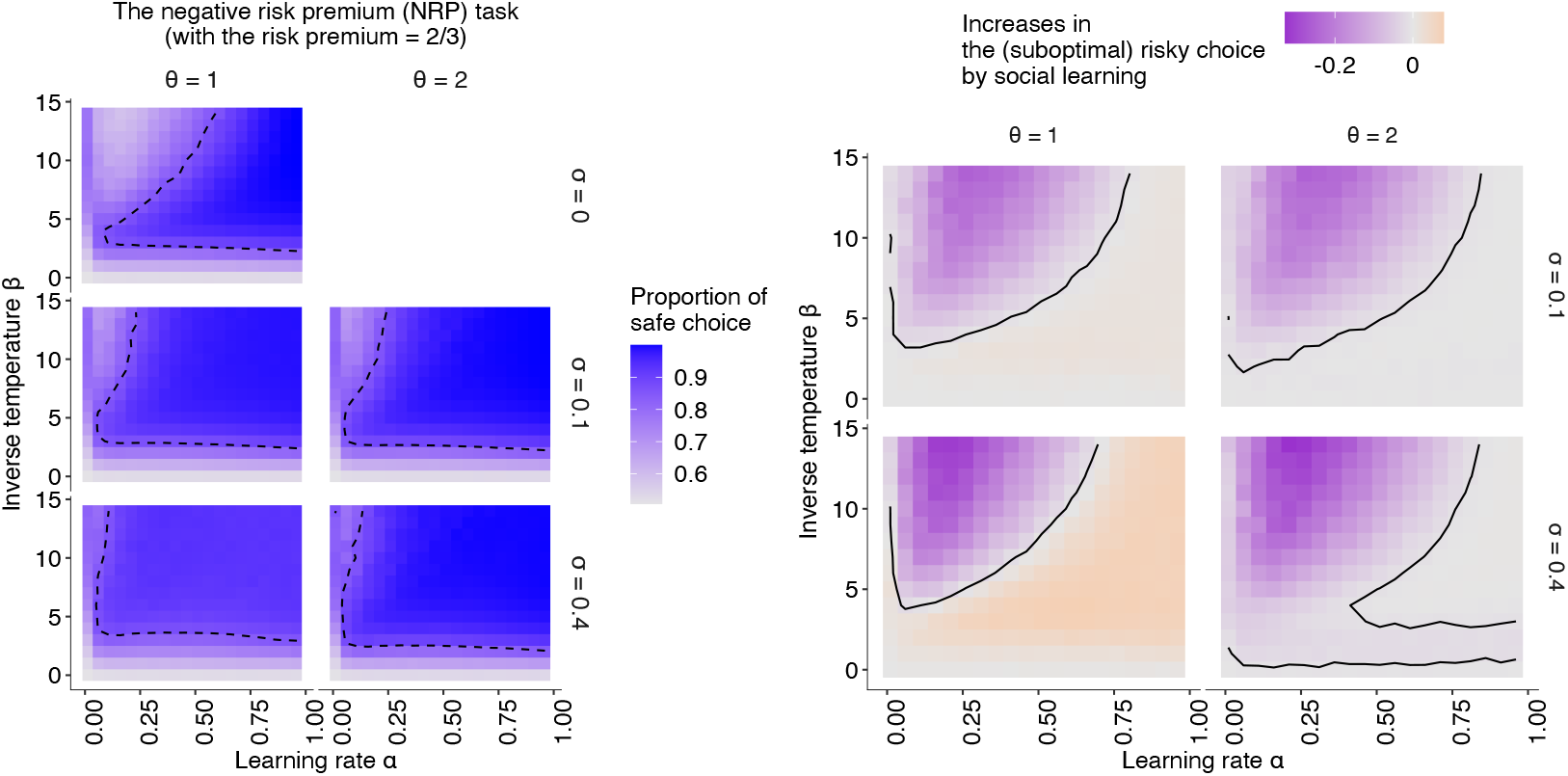
The simulation results under the negative risk premium experimental setup. The relationships between individual learning rate (*α*) and individual inverse temperature (*β*) across different combinations of social learning parameters. (left): The coloured background shows the average proportion of choosing the (optimal) safe option in the second half of the learning trials under social influences with different values of the conformity exponents *θ* and copying weights *σ.* The dashed curve shows the proportion of choosing the safe option at *P_s_* = 0.85. (right): The differences between the mean proportion of risk aversion of asocial learners and that of social learners, highlighting regions in which (suboptimal) risk-seeking increases (orange) and (optimal) risk-aversion increases (purple) by social learning.

**Figure 19.**
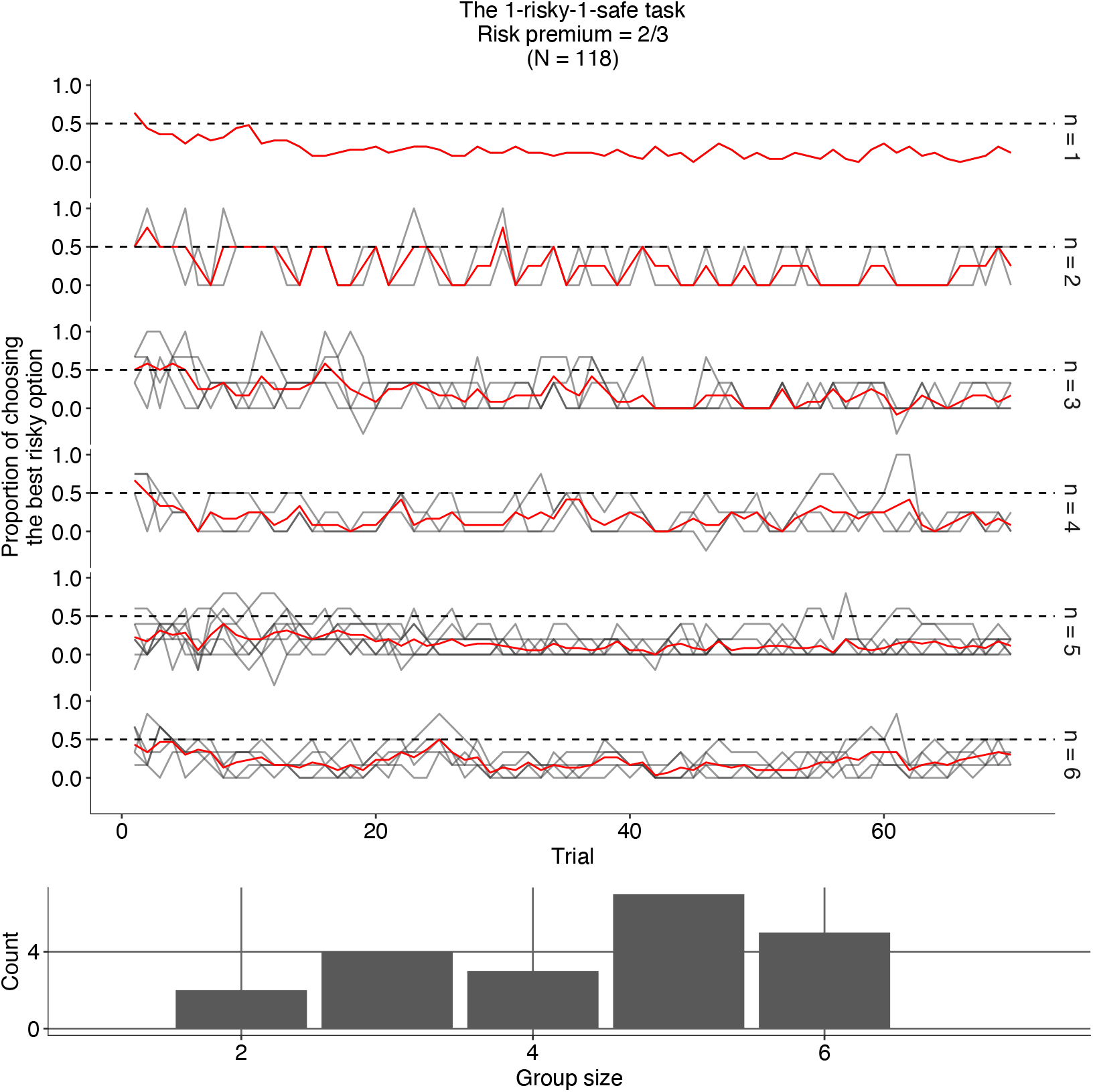
Dynamics of the experimental groups’ mean performance and the distribution of group size. Group size is indicated on the righthand side. Each group’s mean proportion of choosing the best risky option is shown in grey;mean value across the same group-size category is shown in red. The blue line is the mean proportion of choosing the best safe option (that is suboptimal) averaged across the same group-size category.

## Discussion

We have demonstrated that frequency-based copying, one of the most common forms of social learning strategy, can rescue decision makers from committing to adverse risk aversion in a risky trial-and-error learning task, even though a majority of individuals are potentially biased towards suboptimal risk aversion. Although an extremely strong reliance on conformist influence can raise the possibility of getting stuck on a suboptimal option, consistent with the previous view of herding by conformity (***Raafat et al., 2009***; ***Denrell and Le Mens, 2016***), the mitigation of risk aversion and the concomitant collective behavioural rescue could emerge in a wide range of situations under modest use of conformist social learning.

Neither the averaging process of diverse individual inputs nor the speeding up of learning could account for the rescue effect. The individual diversity in the learning rate (*α_i_*) was beneficial for the group performance, whereas that in the social learning weight (*σ_i_*) undermines the average decision performance, which could not be explained simply by a monotonic relationship between diversity and wisdom of crowds (***Lorenz et al., 2011***). Self-organisation through collective behavioural dynamics emerging from the experience-based decision making must be responsible for the seemingly counter-intuitive phenomenon of collective rescue.

Our simplified differential equation model has identified a key mechanism of the collective behavioural rescue: the synergy of positive and negative feedback. Despite conformity, the probability of choosing the suboptimal option can decrease from what is expected by individual learning alone. Indeed, an inherent individual preference for the safe alternative, expressed by the softmax function *e^β_s_Q^*/(*e^βQ_s_^* + *e^βQ_r_^*), is always mitigated by the conformist influence 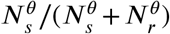 as long as the former is larger than the latter. In other words, risk-aversion was mitigated not because the majority chose the risky option, nor were individuals simply attracted towards the majority. Rather, participants’ choices became risker even though the majority chose the safer alternative at the outset. Intuitively, under social influences (either because of informational or normative motivations), individuals become more explorative, likely to continue samplingthe risky option even after he/she gets disappointed by poor rewards. Once individual risk aversion is reduced, there will exist fewer individuals choosing the suboptimal safe option, which further reduces the number of majority choosing the safe option. This negative feedback facilitates individuals revisiting the risky alternative. Such an attraction to the risky option allows more individuals, including those who are currently sceptical about the value of the risky option, to experience a large bonanza from the risky option, which results in ‘gluing’ them to the risky alternative for a while. Once a majority of individuals get glued to the risky alternative, positive feedback from conformity kicks in, and optimal risk seeking is further strengthened.

Models of conformist social influences have suggested that influences from the majority on individual decision making can lead a group as a whole to collective illusion that individuals learn to prefer any behavioural alternatives supported by many other individuals (***Denrell and Le Mens, 2007***, ***2016***). However, previous empirical studies have repeatedly demonstrated that collective decision making under frequency-based social influences is broadly beneficial and can maintain more flexibility than what suggested by models of herding and collective illusion (***Toyokawa et al., 2019***; ***Aplin et al., 2017***; ***Beckers et al., 1990***; ***Seeley et al., 1991***; ***Camazine et al., 2001***; ***Kandler and Laland, 2013***). For example, ***Aplin et al.*** (***2017***) demonstrated that populations of great tits (*Parus major*) could switch their behavioural tradition after an environmental change even though individual birds were likely to have a strong conformist tendency. A similar phenomenon was also reported in humans (***Toyokawa et al., 2019***).

Although these studies did not focus on risky decision making, and hence individuals were not inherently biased, experimentally induced environmental change was able to create such a situation where a majority of individuals exhibited an out-dated, suboptimal behaviour. However, as we have shown, a collective learning system could rescue their performance even though the individual distribution was strongly biased towards the suboptimal direction at the outset. The great tit and human groups were able to switch their tradition because of, rather than despite, the conformist social influence, thanks to the synergy of negative and positive feedback processes. Such the synergistic interaction between positive and negative feedback could not be predicted by the collective illusion models where individual decision making is determined fully by the majority influence because no negative feedback would be able to operate.

Through online behavioural experiments using a risky multi-armed bandit task, we have confirmed our theoretical prediction that simple frequency-based copying could mitigate risk aversion that many individual learners, especially those who had higher learning rates and/or lower exploration rates, would have exhibited as a result of the hot stove effect. The mitigation of risk aversion was also observed in the NRP task, in which social learning slightly undermined the decision performance. However, because riskiness and expected reward are often positively correlated in a wide range of decision-making environments in the real world (***Pleskac and Hertwig, 2014***), the detrimental effect of reducing optimal risk aversion when risk premium is negative could be negligible in many ecological circumstances, making the conformist social learning beneficial in most cases.

Yet, a majority, albeit a smaller one, still showed risk aversion. The weak re-liance on social learning, which affected only about 15% of decisions, was unable to facilitate strong positive feedback. The little use of social information might have been due to the lack of normative motivations for conformity and to the stationarity of the task. In a stable environment, learners could eventually gather enough information as trials proceeded, which might have made them less curious about information gathering including social learning (***Rendell et al., 2010***). In reality, people might use more sophisticated social learning strategies whereby they change the reliance on social information flexibly over trials (***Deffner et al., 2020***; ***Toyokawa et al., 2017***, ***2019***). Future research should consider more strategic use of social information, and will look at the conditions that elicit heavier reliance on the conformist social learning in humans, such as normative pressures for aligning with majority, volatility in the environment, time pressure, or an increasing number of behavioural options (***Muthukrishna et al., 2015***), coupled with larger group sizes (***Toyokawa et al., 2019***).

The low learning rate *α*, which was at most 0.2 for many individuals in all the experimental task except for the NRP task, should also have hindered the potential benefits of collective rescue in our current experiment, because the benefit of mitigating the hot stove effect would be minimal or hardly realised under such a small susceptibility to the hot stove effect. Although we believe that the simplest stationary environment was a necessary first step in building our understanding of the collective behavioural rescue effect, we would suggest that future studies use a temporally unstable (‘restless’) bandit task to elicit both a higher learning rate and a heavier reliance on social learning, so as to investigate the possibilities of a stronger effect. Indeed, previous studies with changing environments have reported a learning rate as high as *α* > 0.5 (***Toyokawa et al., 2017***, ***2019***; ***Deffner et al., 2020***), under which individual learners should have suffered the hot stove trap more often.

Information about others’ payoffs might also be available in addition to inadvertent social frequency cues in some social contexts (***Bault et al., 2011***), especially with the aid of online communication tools or benevolent pedagogical acts from others. Although communicative acts may transfer information about behavioural alternatives that one has never tried before and may inform about forgone payoffs from other alternatives, which could mitigate the hot stove effect (***Denrell, 2007***; ***Yechiam and Busemeyer, 2006***), it may further amplify the suboptimal decision bias if information senders, despite their cooperative motivation, selectively filter out some pieces of information they think are redundant (***Moussaïd et al., 2015***). In contrast, previous studies suggested that competitions or conflicts of interest among individuals can lead to better collective intelligence than fully cooperative situations (***Conradt et al., 2013***) and can promote adaptive risk taking (***Arbilly et al., 2011***). Further research will identify conditions under which cooperative communication containing richer information can improve decision making and drive adaptive cumulative cultural transmission (***Csibra and Gergely, 2011***; ***Morgan et al., 2015***), when adverse biases in individual decision-making processes prevail.

The generality of our dynamics model should apply to various collective decision-making systems, not only to human groups. Because it is a fundamental property of adaptive reinforcement learning, risk aversion due to the hot stove effect should be widespread in animals (***Real, 1981***; ***Weber et al., 2004***; ***Hertwig and Erev, 2009***). Therefore, its solution, the collective behavioural rescue, should also operate broadly in collective animal decision making because frequency-based copying is one of the common social learning strategies (***Hoppitt and Laland, 2013***; ***Grüter and Leadbeater, 2014***). Future research should determine to what extent the collective behavioural rescue actually impacts animal decision making in wider contexts, and whether it influences the evolution of social learning, information sharing, and the formation of group living.

We have identified a previously overlooked mechanism underlying the adaptive advantages of frequency-based social learning. Our results suggest that an informational benefit of group living could exist well beyond simple informational pooling where individuals can enjoy the wisdom of crowds effect (***Ward and Zahavi, 1973***). Furthermore, the flexibility emerging through the interaction of negative and positive feedback suggests that conformity could evolve in a wider range of environments than previously assumed (***Aoki and Feldman, 2014***; ***Nakahashi et al., 2012***), including temporally variable environments (***Aplin et al., 2017***). Social learning can drive self-organisation, regulating the mitigation and amplification of behavioural biases and canalising the course of repeated decision making under risk and uncertainty.

## Methods

### The approximated dynamics model of collective be-haviour

We assume a group of *N* individuals who exhibit two different behavioural states: choosing a safe alternative *S*, exhibited by *N_S_* individuals;and choosing a risky alternative *R*, exhibited by *N_R_* individuals (*N* = *N_S_* + *N_R_*). We also assume that there are two different ‘inner belief states, labelled ‘−’ and ‘+’. Individuals who possess the negative belief prefer the safe alternative *S* to *R*, while those who possess the positive belief prefer *R* to *S*. A per capita probability of choice shift from one behavioural alternative to the other is denoted by *P*. For example, 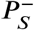 means the individual probability of changing the choice to the safe alternative from the risky alternative under the negative belief. Because there exist 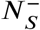 individuals who chose *S* with belief –, the total number of individuals who ‘move on’ to *S* from *R* at one time step is denoted by 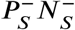. We assume that the probability of shifting to the more preferable option is larger than that of shifting to the less preferable option, that is, 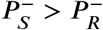 and 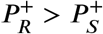 (Fig. 4a).

We assume that the belief state can change by choosing the risky alternative. We define that the per capita probability of becoming + state, that is, having a higher preference for the risky alternative, is *e* (0 ≤ *e* ≤ 1), and hence 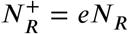. The rest of the individuals who choose the risky alternative become – belief state, that is, 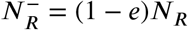.

We define ‘*e*’ so that it can be seen as a risk premium of the gambles. For example, imagine a two-armed bandit task equipped with one risky arm with Gaussian noises and the other a sure arm. The larger the mean expected reward of the risky option (i.e., the higher the risk premium), the more people who choose the risky arm are expected to obtain a larger reward than what the safe alternative would provide. By assuming *e* > 1/2, therefore, it approximates a situation where risk seeking is optimal in the long run.

Here we focus only on the population dynamics: If more people choose *S*, *N_S_* increases. On the other hand, if more people choose *R, N_R_* increases. As a consequence, the system may eventually reach an equilibrium state where both *N_S_* and *N_R_* no longer change. If we find that the equilibrium state of the population (denoted by ★) satisfies 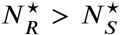, we define that the population exhibits risk seeking, escaping from the hot stove effect. For the sake of simplicity, we assumed 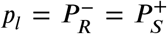 and 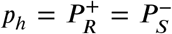, where 0 ≤ *p_l_* ≤ *p_h_* ≤ 1, for the asocial baseline model.

Considering 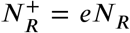 and 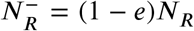, the dynamics are written as the following differential equations:

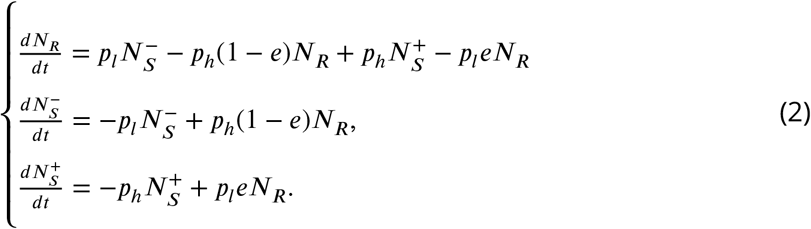

Overall, our model crystallises the asymmetry emerging from adaptive sampling, which is considered as a fundamental mechanism of the hot stove effect (***Denrell, 2007***; ***March, 1996***): Once decision makers underestimate the expected value of the risky alternative, they start avoiding it and do not have another chance to correct the error. In other words, although there would potentially be more individuals who obtain a preference for *R* by choosing the risky alternative (i.e., *e* > 0.5), this asymmetry raised by the adaptive balance between exploration-exploitation may constantly increase the number of people who possess a preference for *S* due to underestimation of the value of the risky alternative. If our model is able to capture this asymmetric dynamics properly, the relationship between *e* (i.e., the potential goodness of the risky option) and *p_l_/p_h_* (i.e., the exploration-exploitation) should account for the hot stove effect, as suggested by previous learning model analysis (***Denrell, 2007***). The equilibrium analysis was conducted in Mathematica ([code is available online](https://github.com/WataruToyokawa/ToyokawaGaissmaier2021/tree/main/dynamicsModel)). The results are shown in Fig. 4.

### Collective dynamics with social influences

For social influences, we assumed that the behavioural transition rates, *P_S_* and *P_R_*, would depend on the number of individuals *N_S_* and *N_R_* as follows:

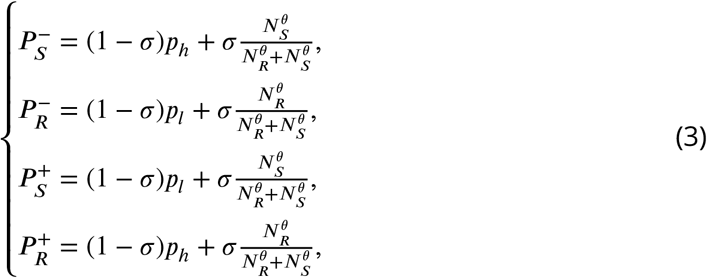

where *σ* is the weight of social influence and *θ* is the strength of the conformist bias, corresponding to the agent-based learning model (Table 1). Other assumptions were the same as in the baseline dynamics model. The baseline dynamics model was a special case of this social influence model with *σ* = 0. Because the system was not analytically tractable, we obtained the numeric solution across different initial distribution of *N*_*S,t*=0_ and *N*_*R,t*=0_ for various combinations of the parameters. The results are shown in Fig. 4d and 5.

### The online experiments

The experimental procedure was approved by the Ethics Committee at the University of Konstanz (‘Collective learning and decision-making study’). Six hundred nineteen English-speaking subjects [294 self-identified as women, 277 as men, 1 as other, and the rest of 47 unspecified;mean (minimum, maximum) age = 35.2 (18, 74) years] participated in the task through the online experimental recruiting platform [Prolific Academic](https://www.prolific.co/). We excluded subjects who disconnected from the online task before completing at least the first 35 rounds from our computational model-fitting analysis, resulting in 585 subjects (the detailed distribution of subjects for each condition is shown in Table 2). A parameter recovery test had suggested that the sample size was sufficient to reliably estimate individual parameters using a hierarchical Bayesian fitting method (see below; Supplementary Fig. 16).

#### Design of the experimental manipulations

The group size was manipulated by randomly assigning different capacities of a ‘waiting lobby’ where subjects had to wait until other subjects arrived. When the lobby capacity was 1, which happened at probability 0.1, the individual condition started upon the first subject’s arrival. Otherwise, the group condition started when there were more than three people at 3 minutes since the lobby opened (see Appendix 1 Supplementary Methods). If there were only two or fewer people in the lobby at this stage, the subjects each were assigned to the individual condition. Note that some groups in the group condition ended up with only two individuals due to a drop out of one individual during the task. The distribution of group size is shown in Supplementary Fig. 17.

We used three different tasks: a 1-risky-1-safe task, a 1-risky-3-safe task, and a 2-risky-2-safe task, where one risky option was expected to give a higher payoff than other options on average (that is, tasks with a positive risk premium (PRP)). To confirm our prediction that risky shift would not strongly emerge when risk premium was negative (i.e. risk seeking was suboptimal), we also conducted another 1-risky-1-safe task with a negative risk premium (the NRP task). Partic-ipants’ goal was to gather as many individual payoff as possible, as monetary incentives were given to the individual performance. In the NRP task, risk aversion was favourable instead. All tasks had 70 decision-making trials. The task proceeded on a trial basis;that is, trials of all individuals in a group were synchro-nised. Subjects in the group condition could see social frequency information, namely, how many people chose each alternative in the preceding trial. No social information was available in the first trial. These tasks were assigned randomly as a between subject condition, and subjects were allowed to participate in one session only.

We employed a skewed payoff probability distribution rather than a normal distribution forthe risky alternative, and we conducted not only a two-armed task but also four-armed bandit tasks, because our pilot study had suggested that subjects tended to have a small susceptibility to the effect (*α_i_*(*β_i_* + 1) ≪ 2), and hence we needed more difficult settings than the conventional Gaussian noise binary-choice task to elicit risk aversion from individual decision makers. Running agentbased simulations, we confirmed that these task setups used in the experiment could elicit the collective rescue effect (Supplementary Fig. 11 and 18).

The details of the task setups are as follows:

##### The 1-risky-1-safe task

The optimal risky option produced either 50 or 550 points at probability 0.7 and 0.3, respectively (the expected payoff was 200). The safe option produced 150 points (with a small amount of Gaussian noise with s.d. = 5).

##### The 1-risky-3-safe task

The optimal risky option produced either 50 or 425 points at probability 0.6 and 0.4, respectively (the expected payoff was 200). The three safe options each produced 150, 125, and 100 points, respectively, with a small Gaussian noise with s.d. = 5.

##### The 2-risky-2-safe task

The optimal risky option produced either 50 or 425 points at probability 0.6 and 0.4, respectively (the expected payoff was 200). The two safe options each produced 150 and 125 points, respectively, with a small Gaussian noise with s.d. = 5. The suboptimal risky option, whose expected value was 125, produced either 50 or 238 points at probability 0.6 and 0.4, respectively.

##### The negative risk premium task

The setting was the same as in the 1-risky-1-safe task, except that the expected payoff from the risky option was smaller than the safe option, producing either 50 or 220 points at probability 0.7 and 0.3, respectively (the expected payoff was 101).

We have confirmed through agent-based model simulations that the collective behavioural rescue could emerge in tasks equipped with the experimental settings (Supplementary Fig. 11). We have also confirmed that risk seeking does not always increase when risk premium is negative (Supplementary Fig. 18). With the four-armed tasks we aimed to demonstrate that the rescue effect is not limited to binary-choice situations. Other procedures of the collective learning task were the same as those used in our agent-based simulation shown in the main text. The experimental materials including illustrated instructions can be found in Supplementary Video 1 (individual condition) and Video 2 (group condition).

### The hierarchical Bayesian model fitting

To fit the learning model, we used a hierarchical Bayesian method, estimating the global means (*μ_α_*, *μ_β_*, *μ_σ_*, and *μ_θ_*) and global variances (*v_α_*, *ν_β_*, *ν_σ_*, and *ν_θ_*) for each of the three experimental conditions and forthe individual and group conditions separately. For the individual condition, we assumed *σ* = 0 for all subjects and hence no social learning parameters were estimated. Full details of the modelfitting procedure and prior assumptions are shown in the Supplementary Methods. The R and Stan code used in the model fitting are available from an online repository.

#### Parameter recovery and post hoc simulation

To assess the adequacy of the hierarchical Bayesian model-fitting method, we tested how well the hierarchical Bayesian method (HBM) could recover ‘true’ parameter values that were used to simulate synthetic data. We simulated artificial agents’ behaviour assuming that they behave according to the social learning model with each parameter setting. We generated ‘true’ parameter values for each simulated agent based on both experimentally fit global parameters (Table 1; parameter recovery test 1). In addition, we ran another recovery test using arbitrary global parameters that deviated from the experimentally fit values (parameter recovery test 2), to confirm that our fitting procedure was not just ‘attracted’ to the fit value. We then simulated synthetic behavioural data and recovered their parameter values using the HBM described above. Both parameter recovery tests showed that all the recovered individual parameters were positively correlated with the true values, whose correlation coefficients were all larger than 0.5. We also confirmed that 30 of 32 global parameters in total were recovered within the 95% Bayesian credible intervals, and that even those two non-recovered parameters (*μ_β_* for the 2-risky-2-safe task in parameter recovery test 1 and *μ_α_* for the 1-risky-3-safe task in parameter recovery test 2) did not deviate so much from the true value (Supplementary Fig. 16).

We also ran a series of individual-based model simulations using the calibrated global parameter values for each condition. First, we randomly sampled a set of agents whose individual parameter values were drawn from the fit global parameters. Second, we let this synthetic group of agents perform the task for 70 rounds. We repeated these steps 100,000 times for each task setting and for each individual and group condition.

## Appendix 1: Supplementary methods

### An analytical result derived by Denrell (2007)

In the simplest setup of the two-armed bandit task, ***Denrell*** (***2007***) derived an explicit form for the asymptotic probability of choosing the risky alternative 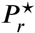 (as *t* → ∞) as follows:

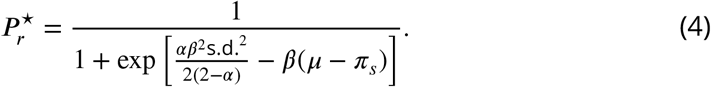

Equation 4 identifies a condition under which reinforcement learners exhibit risk aversion. In fact, when there is no risk premium (i.e., *μ* ≤ *π_s_*), the condition of risk aversion always holds, that is, 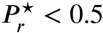. Consider the case where risk aversion is suboptimal, that is, *μ* > *π_s_*. Equation 4 suggests that suboptimal risk aversion emerges when learning is myopic (i.e., when *α* is large) and/or decision making is less explorative (i.e., when *β* is large). For instance, when the payoff distribution of the risky alternative is set to *μ* = *π_s_* + 0.5 and s.d.^2^ = 1, the condition of risk aversion, 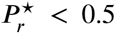, holds under *β* > (2 – *α*)/*α*, which corresponds to the area above the dashed curve in Fig. 1b in the main text. Risk aversion becomes more prominent when the risk premium *μ* – *π_s_* is small and/or the payoff variance s.d.^2^ is large.

### The online experiments

#### Subjects

The positive risk premium (PRP) tasks were conducted between August and October 2020 (recruiting 492 subjects), while the negative risk premium (NRP) task was conducted in September 2021 (recruiring 127 subjects) in response to the comments from peer reviewers. All subjects declared their residence in the United Kingdom, the United States, Ireland, or Australia. All subjects consented to participation through an online consent form at the beginning of the task. We excluded subjects who disconnected from the online task before completing at least the first 35 rounds from our computational model-fitting analysis, resulting in 467 subjects for the PRP tasks and 118 subjects for the NRP task (the detailed distribution of subjects for each condition is shown in Table 1 in the main text). The task was available only for English-speaking subjects and they had to be 18 years old or older. Only subjects who passed a comprehension quiz at the end of the instructions could enter the task. Subjects were paid 0.8 GBP as a show-up fee as well as an additional bonus payment depending on their performance in the decision-making task In the PRP tasks 500 artificial points were converted to 8 pence, while in the NRP task 500 points were converted to 10 pence so as to compensate the less productive environment, resulting in a bonus ranging between £1.0 and £3.5.

#### Sample size

Our original target sample size for the PRP tasks was 50 subjects for the individual condition and 150 subjects for the group condition where our target average group size was 5 individuals per group. For the NRP task, we aimed to recruit 30 individuals for the individual condition and 100 individuals (that is, 20 groups of 5) for the group condition. Subjects each completed 70 trials of the task. The sample size and the trial number had been justified by a model recovery analysis of a previous study (***Toyokawa et al., 2019***).

Because of the nature of the ‘waiting lobby’, which was available only for 3 minutes, we could not fully control the exact size of each experimental group. Therefore, we set the maximum capacity of a lobby to 8 individuals for the 1-safe-1-risky task, which was conducted in August 2020, so as to buffer potential dropouts during the waiting period. Since we learnt that dropping out happened far less than we originally expected (Supplementary Fig. 17), we reduced the lobby capacity to 6 for both the 1-risky-3-safe and the 2-risky-2-safe task, which were conducted in October 2020. As a result, we had 20 groups (mean group size = 6.95), 21 groups (mean group size = 4.7), 19 groups (mean group size = 4.3), and 21 gorups (mean group size = 4.4), for the 1-risky-1-safe, 1-risky-3-safe, 2-risky-2-safe task, and the negative risk premium 2-armed task, respectively. Although we could not achieve the sample size targeted, partly due to the dropouts during the task and to a fatal error occurring in the experimental server in the first few sessions of the four-armed tasks, the parameter recovery test with *N* = 105 suggested that the current sample size should be reliable enough to estimate social influences for each subject (Supplementary Fig. 16).

### The hierarchical Bayesian parameter estimation

We used the hierarchical Bayesian method (HBM)to estimate the free parameters of our learning model. HBM allowed us to estimate individual differences, while this individual variation is bounded by the group-level (i.e., hyper) parameters. To do so, we used the following non-centred reparameterisation (the ‘Matt trick’) as follows:

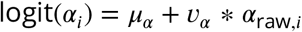

where *μ_α_* is a global mean of logit(*α_i_*) and *ν_α_* is a global scale parameter of the individual variations, which is multiplied by a standardised individual random variable *α*_raw,*i*_. We used a standardised normal prior distribution centred on 0 for *μ_α_* and an exponential prior for *v_α_*. The same method was applied to the other learning parameters *β_i_, σ_i_*, and *θ_i_.*

We assumed that the ‘raw’ values of individual random variables (*α*_raw,*i*_, *β*_raw,*i*_, *σ*_raw*i*_, *θ*_raw,*i*_) were drawn from a multivariate normal distribution. The correlation matrix was estimated using a Cholesky decomposition with a weakly informative Lewandowski-Kurowicka-Joe prior that gave a low likelihood to very high or very low correlations between the parameters (***McElreath, 2020***; ***Deffner et al., 2020***).

#### Model fitting

All models were fitted using the Hamiltonian Monte Carlo engine CmdStan 2.25.0 (https://mc-stan.org/cmdstanr/index.html) in R 4.0.2 (https://www.r-project.org). The models contained at least six parallel chains and we con-firmed convergence of the Markov chain Monte Carlo (MCMC) using both the Gelman-Rubin statistics criterion 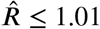 and the effective sample sizes greater than 500. The R and Stan code used in the model fitting are available from an online repository.

### The value-shaping social influence model

We considered another implementation of social influences in reinforcement learning, namely, a value-shaping (***Najar et al., 2020***) (or ‘outcome-bonus’ (***Biele et al., 2011***)) model rather than the decision-biasing process assumed in our main analyses. In the value-shaping model, social influence modifies the *Q* value’s up-dating process as follows:

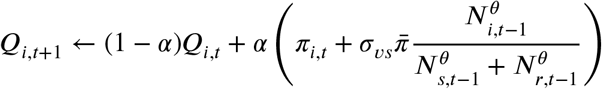

where the social frequency cue acts as an additional ‘bonus’ to the value that was weighted by *σ_vs_* (*σ_vs_* > 0)and standardised by the expected payoff from choosing randomly among all alternatives 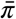. Here we assumed no direct social influence on the action selection process (i.e., *σ* = 0 in our main model). We confirmed that the collective behavioural rescue could emerge when the inverse temperature *β* was sufficiently small (Supplementary Fig. 8). Although it is beyond the focus of this article whether people’s behaviour can be fit better by this value-shaping model than by the decision-biasing model, it is an interesting question for future research (***Najar et al., 2020***).

## Appendix 2: Supplementary videos

**Supplementary Video 1. A sample screenshot of the online experimental task (Individual condition).** This video was taken only for the demonstration purpose and hence not associated to any actual participant’s behaviour.

**Supplementary Video 2. A sample screenshot of the online experimental task with N = 3 (group condition).** This video was taken only for the demonstration purpose and hence not associated to any actual participant’s behaviour. Also note that actual participants could see only one browser window per participant in the experimental sessions.

## Appendix 3: Supplementary figures

**Figure 7. (Figure 1 - figure supplement 1).**
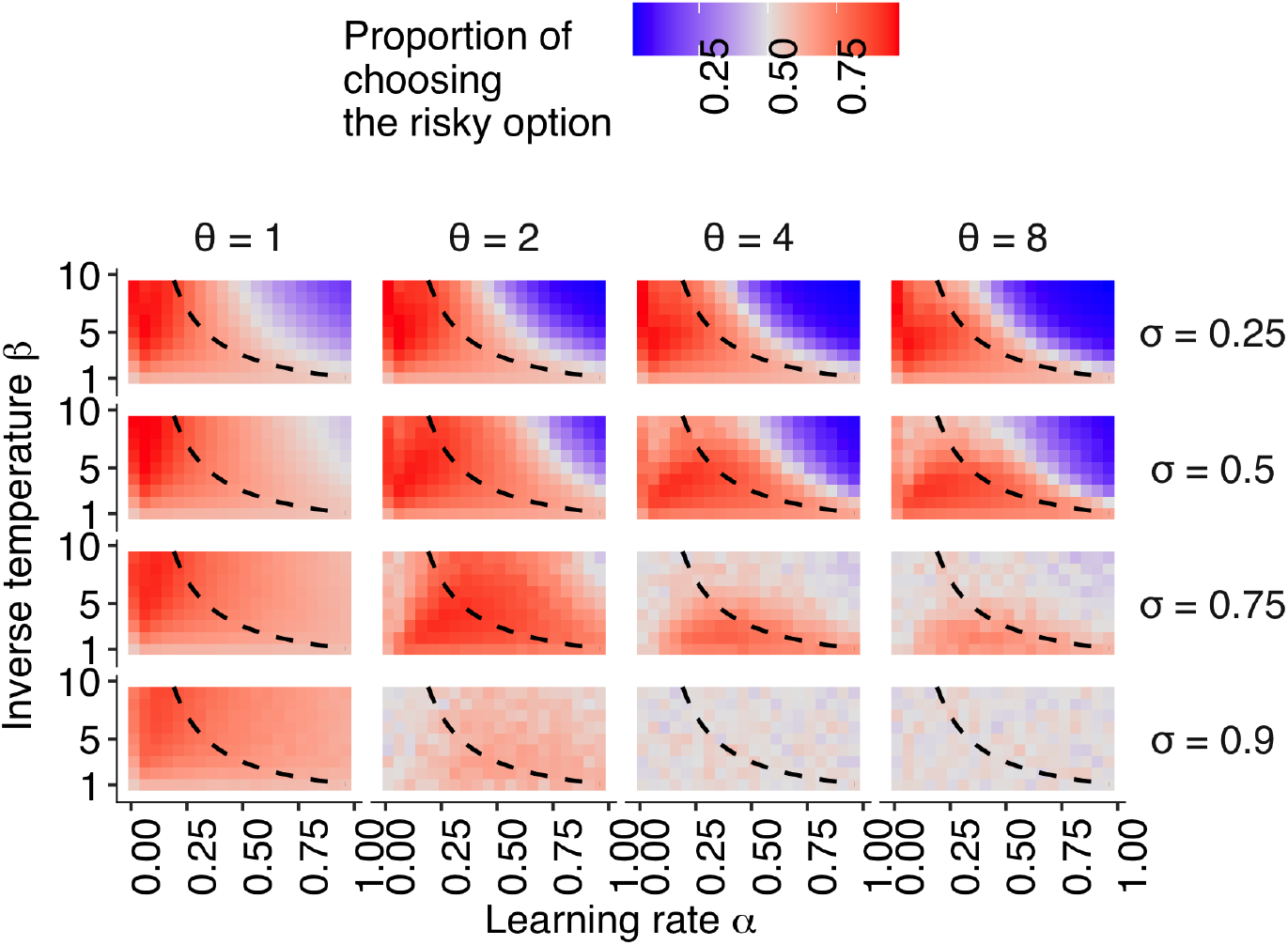
The emergence of risk-taking bias due to the hot stove effect depending on a combination of the reinforcement learning parameters. The effect of the relationship between individual learning rate (*α*) and individual inverse temperature (*β*) across the different combinations of social learning parameters on the mean proportion of choosing the risky alternative in the second half of the trials of the two-armed bandit task described in Fig. 1 in the main text. The dashed curves give a set of parameter combinations with which asocial learners are expected to choose the risky alternative in the same proportion as they choose the safe alternative (i.e., 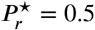) in the infinite time horizon *T* → ∞, given by *β* = (2 – *α*)/*α*.

**Figure 8. (Figure 1 - figure supplement 2).**
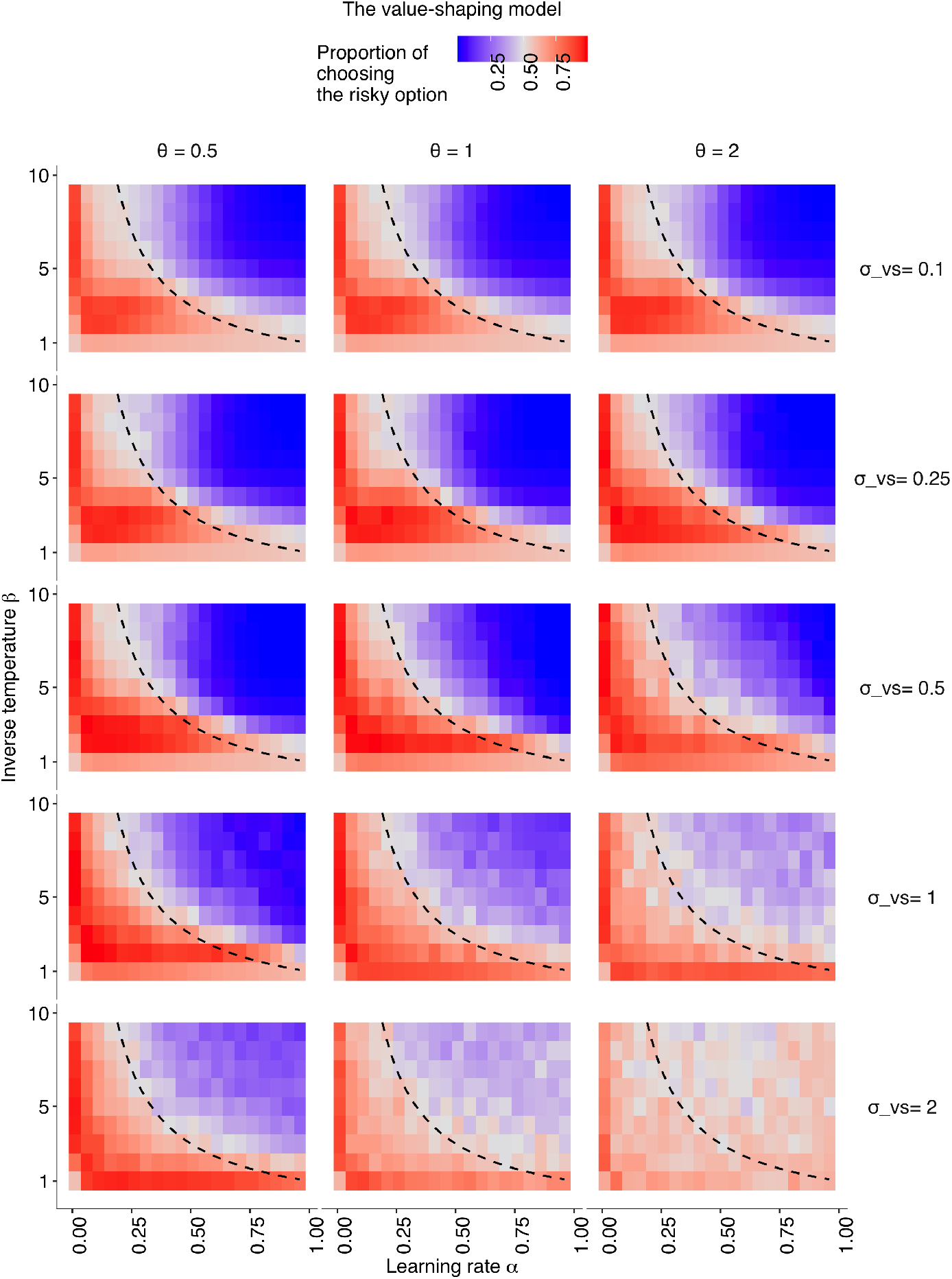
The results of the value-shaping social influence model. The relationships between individual learning rate (*α*) and individual inverse temperature (*β*) across different combinations of social learning parameters. The coloured background shows the average proportion of choosing the risky option in the second half of the learning trials *P*_*r,t*>75_ > 0.5. Different social learning weights (*σ_vs_*) are shown from top to bottom (*σ_vs_* ∈ {0,0.1,0.25,0.5,1,2}). Different conformity exponents are shown from left to right (*θ* ∈ {0.5, 1,2}). The dashed curve is the asymptotic equilibrium at which asocial learners are expected to end up choosing both alternatives with equal likelihood (i.e., 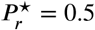), given by *β* = (2 – *α*)/*α*.

**Figure 9. (Figure 1 - figure supplement 3).**
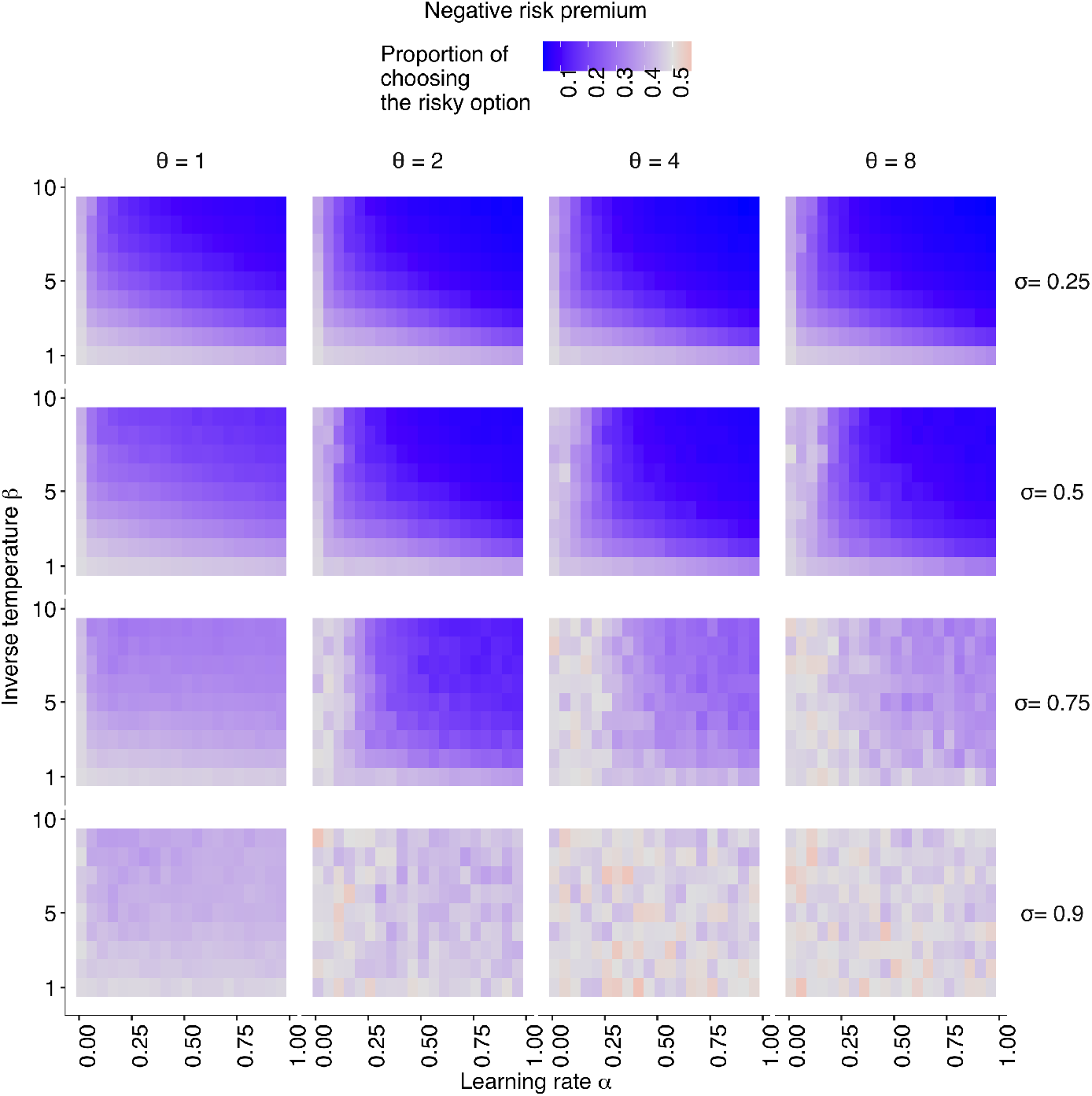
The simulation result with the negative risk premium. The relationships between individual learning rate (*α*) and individual inverse temperature (*β*) across different combinations of social learning parameters. The coloured background shows the average proportion of choosing the risky option in the second half of the learning trials *P*_*r,t*>75_ > 0.5. Different social learning weights (*σ*) are shown from top to bottom (*σ* ∈ {0,0.25,0.5,0.75,0.9}). Different conformity exponents are shown from left to right (*θ* ∈ {1,2,4,8}). The risk premium is negative *μ* = −0.5.

**Figure 10. (Figure 1 - figure supplement 4).**
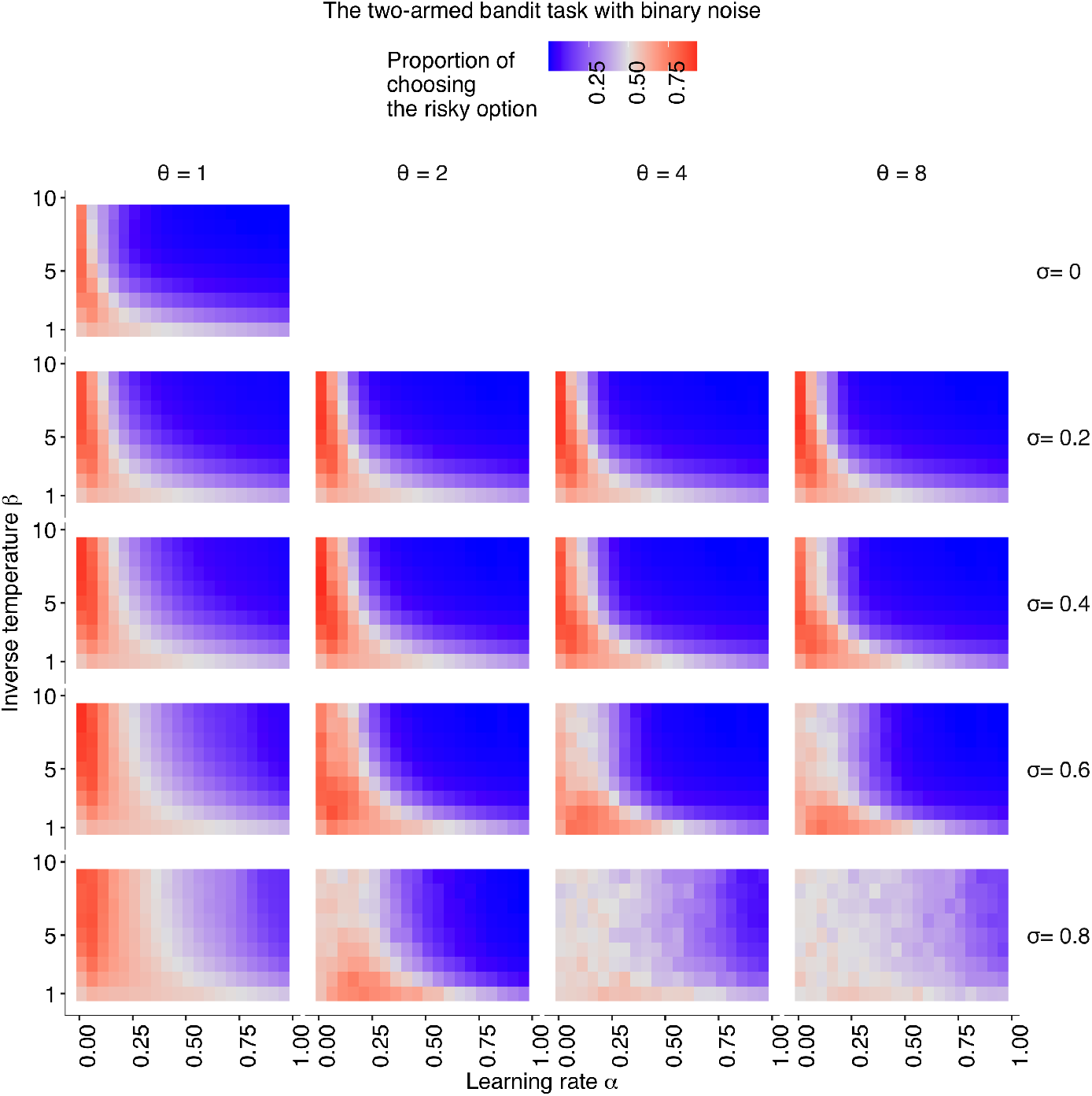
The simulation result with the Bernoulli noise distribution. The relationships between individual learning rate (*α*) and individual inverse temperature (*β*) across different combinations of social learning parameters. The coloured background shows the average proportion of choosing the risky option in the second half of the learning trials *P*_*r,t*>75_ > 0.5. Different social learning weights (*σ*) are shown from top to bottom (*σ* ∈ {0, 0.2, 0.4, 0.6, 0.8}). Different conformity exponents are shown from left to right (*θ* ∈ {1,2,4,8}). The binary payoff distribution was used where the safe alternative always provides *π_s_* = 1 while the risky alternative provides either a 70% chance of *π_r_* = 0 or a 30% chance of *π_r_* = 5. The risk premium was 1.5.

**Figure 11. (Figure 1 - figure supplement 5).**
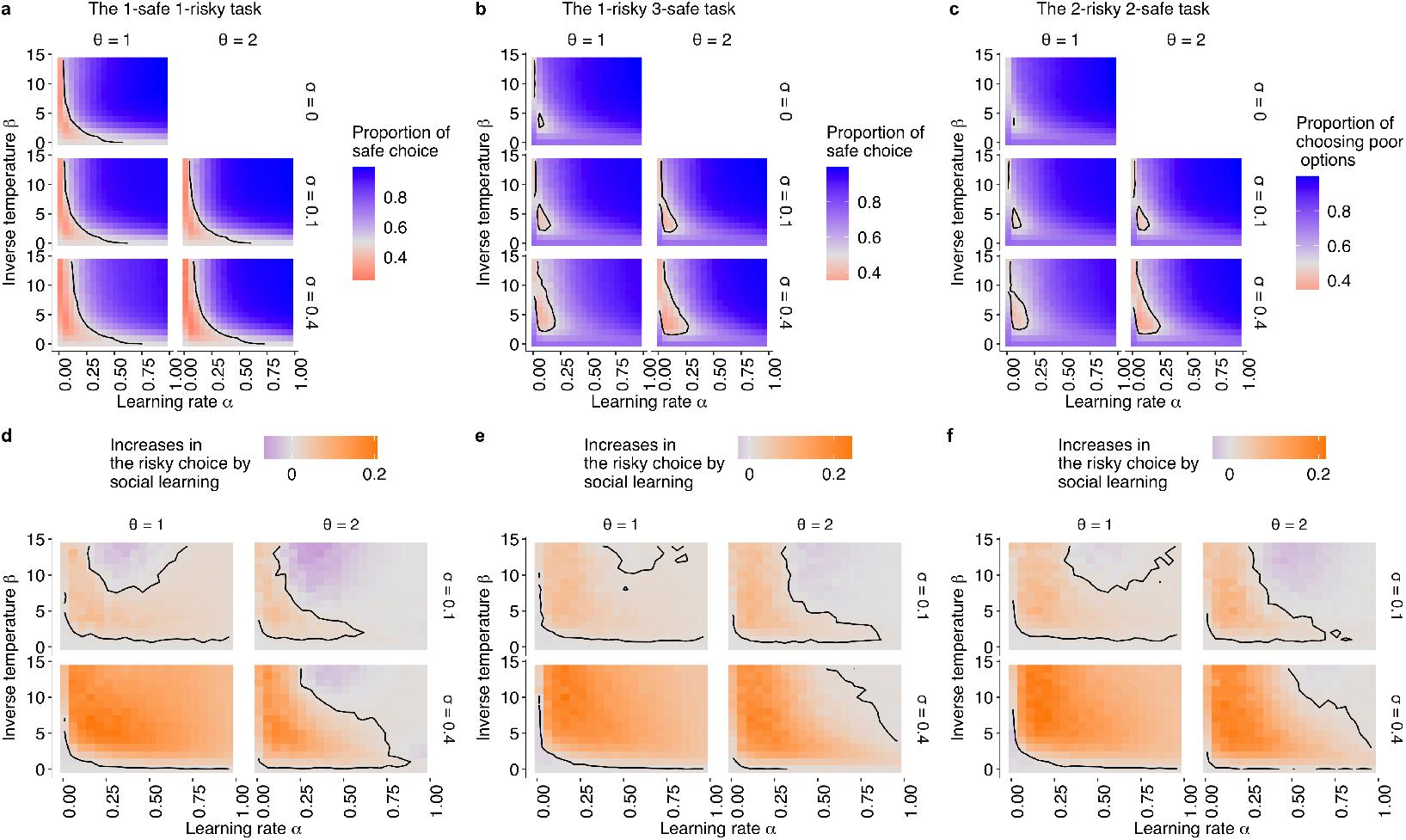
The simulation results under the positive risk premium experimental setups (a,d: the 1-risky-1-safe;b,e: the 1-risky-3-safe;c,f: the 2-risky-2-safe). The relationships between individual learning rate (*α*) and individual inverse temperature (*β*) across different combinations of social learning parameters. (a-c): The coloured background shows the average proportion of choosing the risky option in the second half of the learning trials (*P*_*r,t*>75_ > 0.5) under social influences with different values of the conformity exponents *θ* and copying weights *σ*. The dashed curve is the asymptotic equilibrium at which asocial learners are expected to end up choosing the two alternatives with equal likelihood (i.e., *P_r_* = 0.5). (d-f): The differences between the mean proportion of risk aversion of asocial learners and that of social learners, highlighting regions in which performance is improved (that is, risk-seeking increases; orange) or undermined (that is, risk-aversion is amplified;purple) by social learning.

**Figure 12. (Figure 2 - figure supplement 1).**
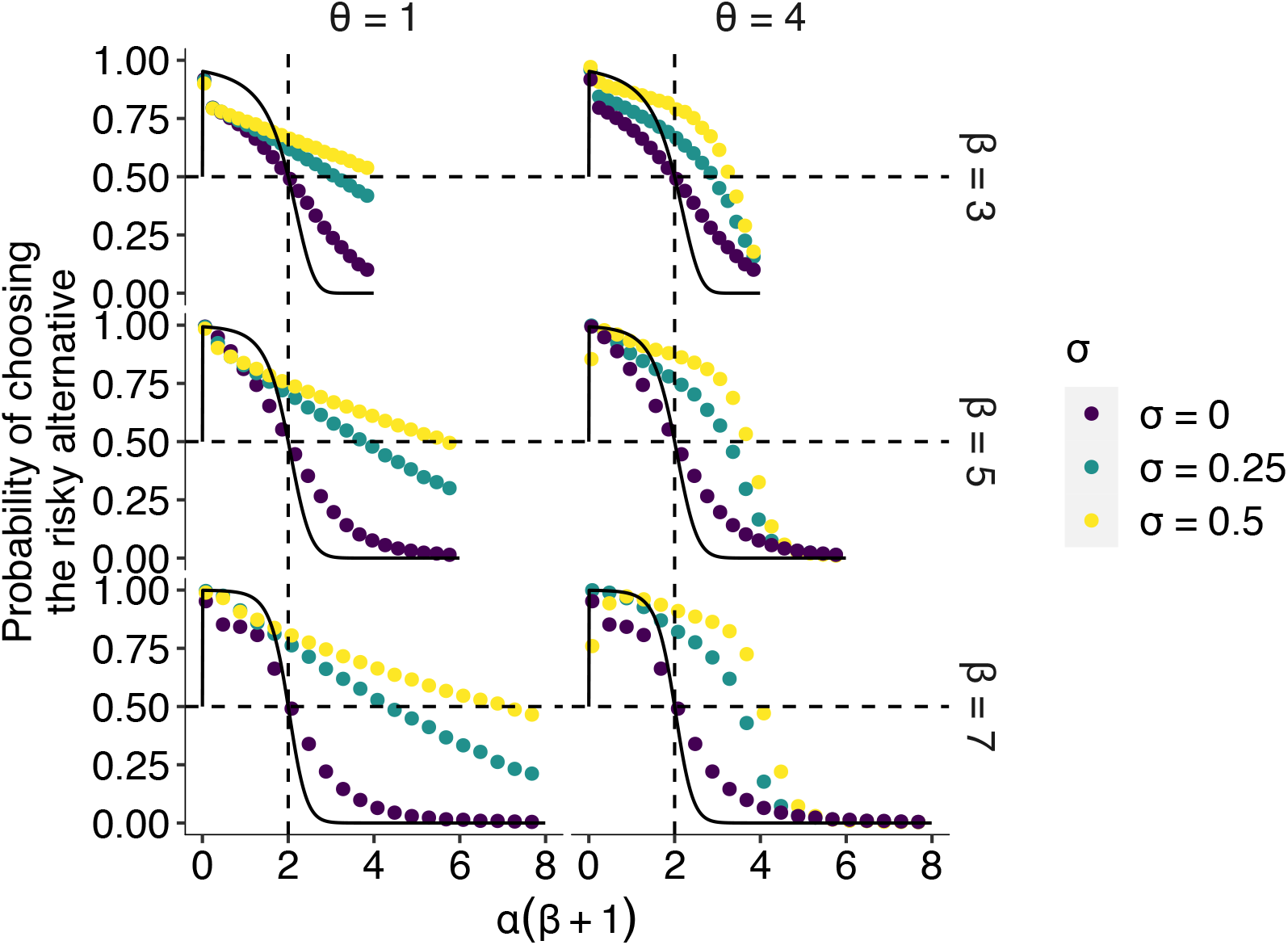
The effect of social learning on the average decision performance on the longer time horizon. The *x* axis is an interaction of two reinforcement learning parameters *α*(*β* + 1), that is, the susceptibility to the hot stove effect. The *y* axis is the mean probability of choosing the optimal risky alternative in the last 75 trials in the two-armed bandit task whose setup was the same as in Figs. 1 and 2 in the main text (i.e., *μ* = 0.5, s.d. = 1) except for the longer time horizon *T* = 1075 compared to the time horizon used in the main text (*T* = 150). The dotted curves are the mean result of agent-based simulations of groups of social learners with two different mean values of the copying weight *σ* ∈ {0.25,0.5} or individual learners with *σ* = 0. Each panel shows a different combination of the inverse temperature *β* and the conformity exponent *θ*. The black solid curve is the theoretical benchmark where individual reinforcement learners were expected to asymptote with *T* → +∞. Compared to Fig. 2 in the main text, individual learners got closer to the benchmark. On the other hand, the performance of social learners remained deviated from the benchmark, suggesting that social influence had a qualitative impact on the course of learning and decision making, rather than merely slowing down approaching the equilibrium of individual learning.

**Figure 13. (Figure 2 - figure supplement 2).**
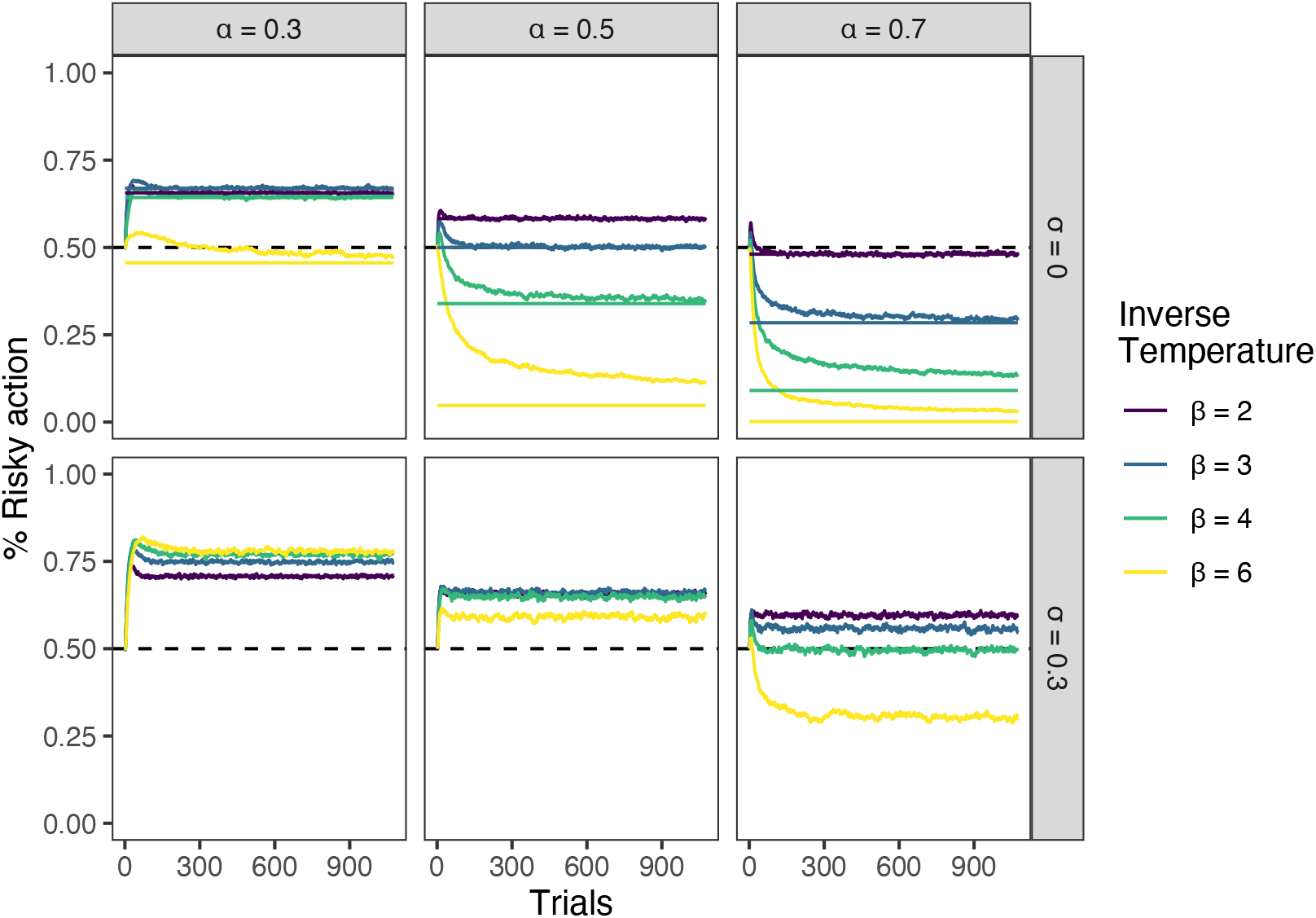
The effect of social learning on the time evolution of decision performance. The *x* axis is the number of trials. The *y* axis is the mean proportion of choosing the optimal risky alternative. Each colour shows a different *β*. For the asocial learning condition (i.e., *σ* = 0), the analytical benchmark to which reinforcement learners asymptote is shown as a horizontal line. Conformity exponent *θ* was 2. Group size was 8. The simulation was repeated 1,000 times for each combination of parameters. Compared to asocial learning cases, social learning (*σ* = 0.3) qualitatively alters the course of learning, rather than just speeding up or slowing down learning.

**Figure 14. (Figure 4 - figure supplement 1).**
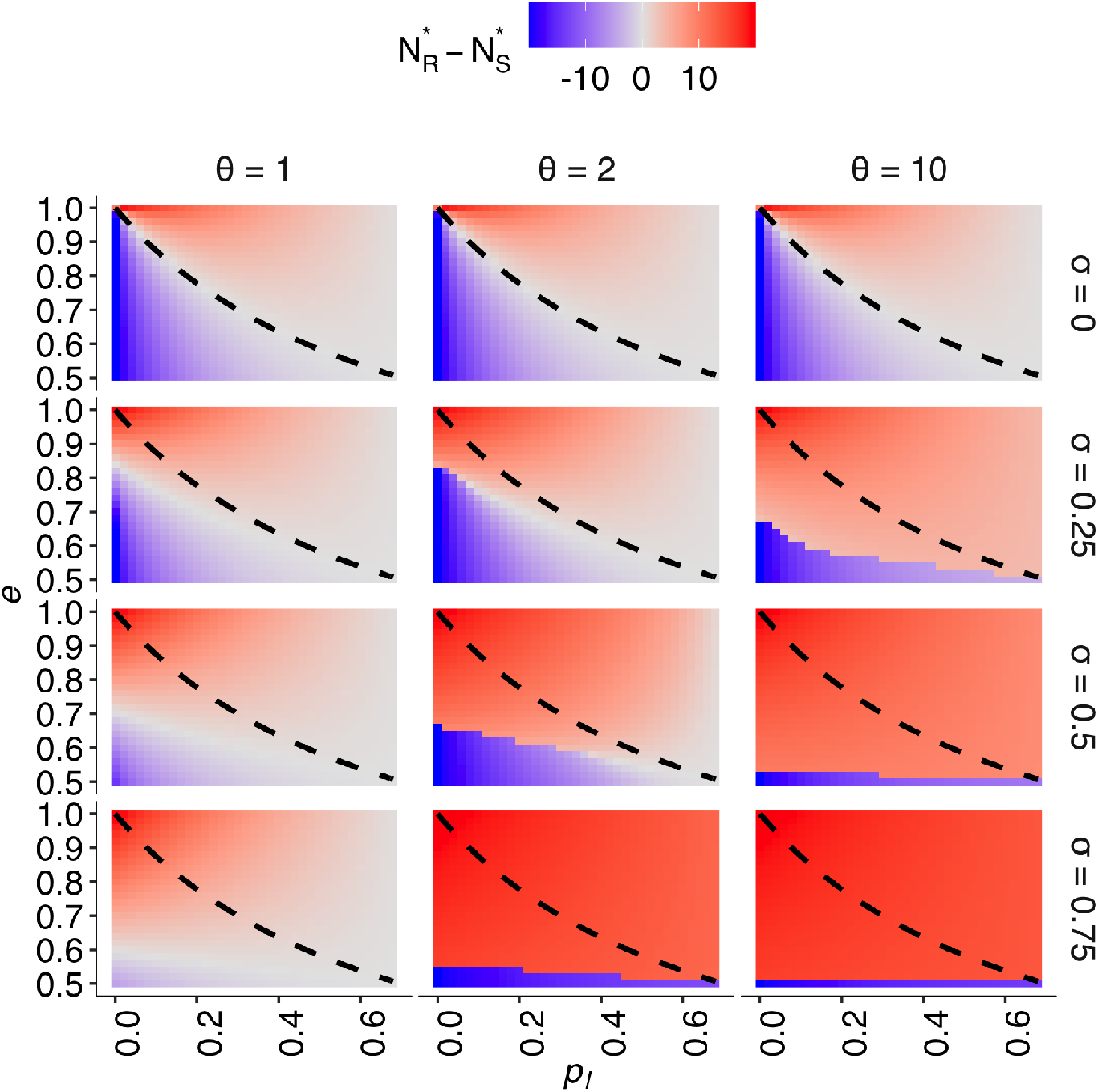
The result of the differential equation model. The effect of both the per capita probability of exploration *p_l_* and *e* (i.e., the ratio of individuals who prefer behavioural state *R*) on the equilibrium degree of risk seeking (i.e., 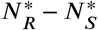), across the different combinations of social influence parameters. Different social influence weights are shown from top to bottom (*σ* ∈ {0,0.25,0.5,0.75}). Different conformity exponents are shown from left to right (*θ* ∈ {1,2,10}). The dashed curve is *e* = *p_h_*/(*p_h_* + *p_t_*). The numeric solution was obtained with conditions 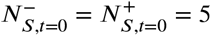, *N_R,t_*=0 = 10, and *p_h_* = 0.7.

**Figure 15. (Figure 5 - figure supplement 1).**
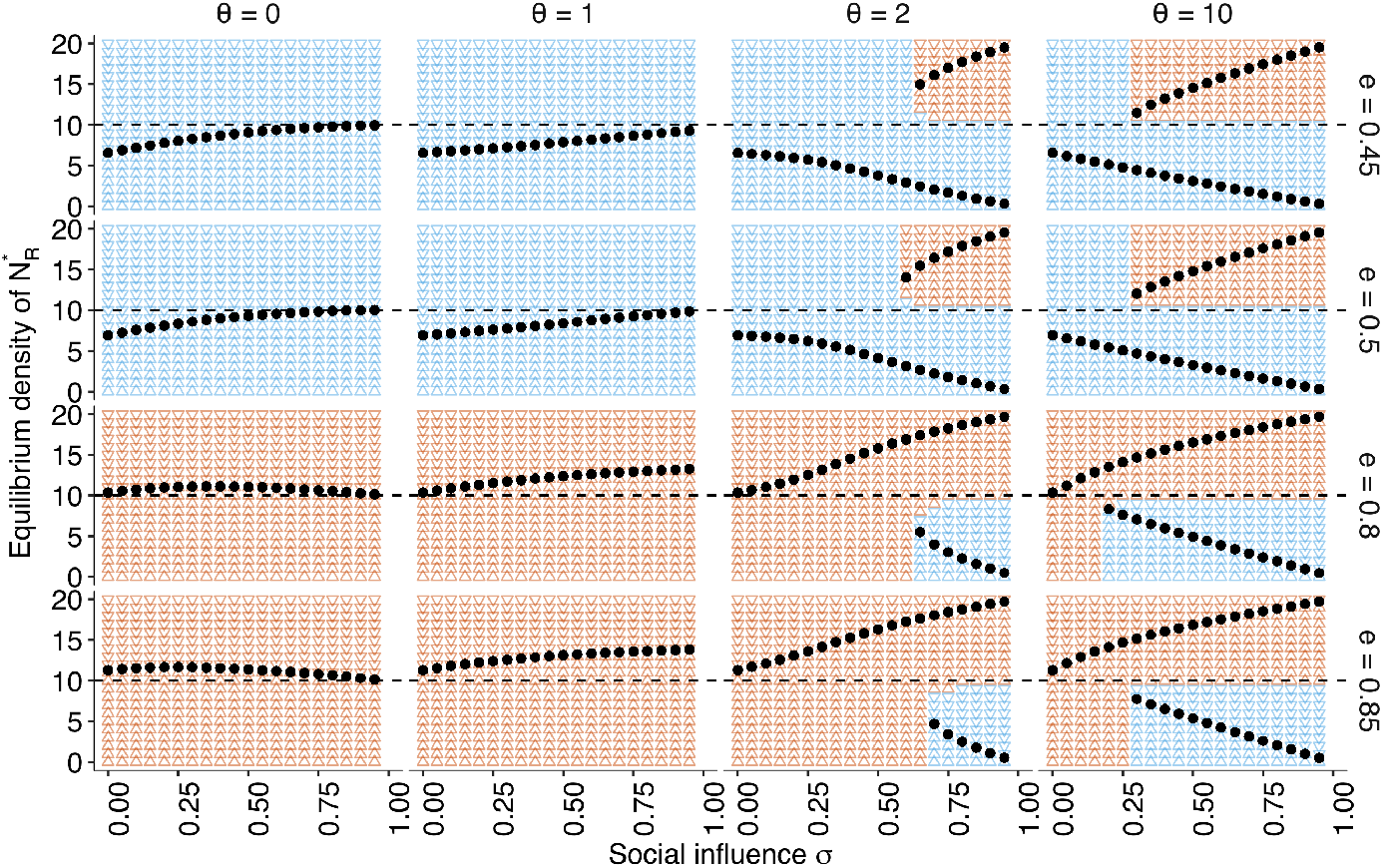
The approximate bifurcation analysis. The relationship between the social influence weight *σ* and the equilibrium number of individuals choosing the risky alternative 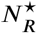 across the different conformity exponents *θ*(∈ {0,1,2,10}), shown as black dots. The triangular points shown in the background of each panel indicate regions in which the group approaches risk aversion (i.e., 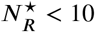; blue) or the risk-seeking equilibrium (i.e., 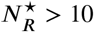; red). Two different equilibria mean that the system has a bifurcation under a given *σ*. The direction of the background triangles indicates whether *N_R_* increases (Δ) or decreases (∇) relative to its starting position. The other parameters are set to *p_h_* = 0.7, *p_l_* = 0.2.

**Figure 16.**
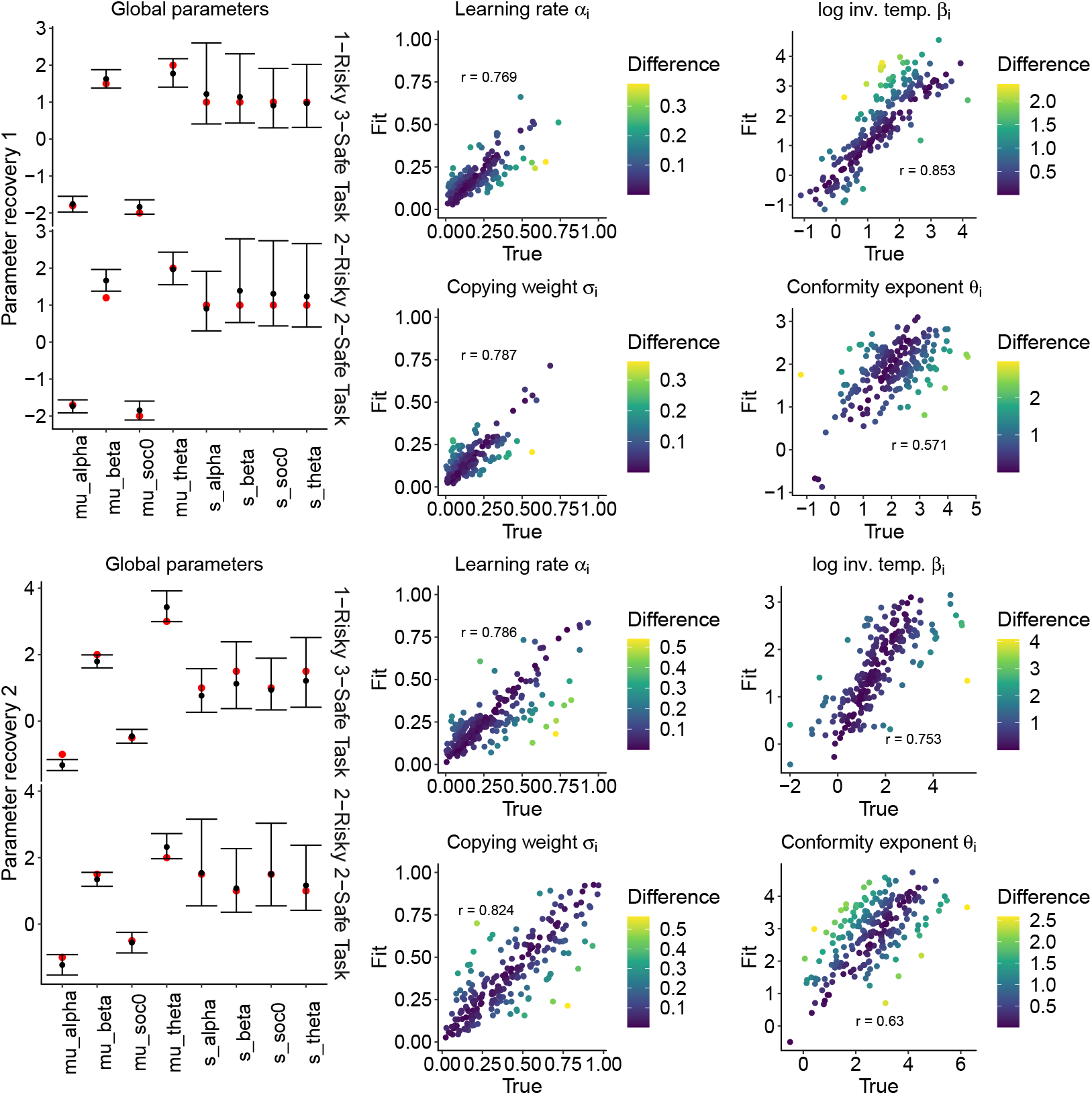
The parameter recovery performance. The top half and bottom half of the figure are the results of parameter recovery test 1 and 2, respectively. The left column shows the global parameters fitted for each of the two four-armed bandit tasks, the 1-risky-3-safe task (*N* = 105) and the 2-risky-2-safe task (*N* = 105). The red points are the true values and the black points are the mean posterior values (i.e., recovered values). The 95% Bayesian credible intervals are shown with error bars. The middle and right column are individual-level parameters across the two task conditions (*N* = 210). The *x* axis is the true value and the*y* axis is the fitted (i.e., the mean posterior) individual value. The differences between the true value and the estimated value are shown in different colours (Dark: fit well). The Pearson’s correlation coefficients between the true and fitted values are shown.

## Data availability

Both experimental and simulation data are available in an online repository: https://github.com/WataruToyokawa/ToyokawaGaissmaier2021

## Acknowledgements

This work was funded by a Small Project Grant from the Centre for the Advanced Study of Collective Behaviour, the University of Konstanz (S20-06), by the University of Konstanz Committee on Research (FP031/19), and by the Deutsche Forschungsgemeinschaft (DFG, German Research Foundation) under Germany’s Excellence Strategy - EXC 2117 - 422037984. We thank Iain Couzin, Lucy Aplin, Brendan Barrett, Ralf Kurvers, Charley Wu, and Anita Todd for many helpful comments on earlier versions of this paper.

## Author contributions

W.T. planned the study, built and analysed the mathematical model, made the experimental material, and ran the online experiment. W.T. and W.G. analysed the data, discussed and refined the experimental design, and wrote the manuscript.

## Materials & Correspondence

Correspondence and requests for materials should be addressed to W.T.

## Notes

https://github.com/WataruToyokawa/ToyokawaGaissmaier2021

### Competing Interest Statement

The authors have declared no competing interest.

### Summary of Updates

We have revised the manuscript so that relationships between our model and the previous literature are highlighted in detail, specifying how our modelling serves as a crucial extension of the previous approach to disentangle the puzzling relationship between maladaptive herding (or collective illusion) and collective intelligence under systematically biased individual decision making. We have also added additional behavioural experimental results to explore a wider range of environments and have added discussion about implications of the differences between informational and normative motivations of conformity.

